# Beyond Coral Cover A framework for assessing the condition of coral reef habitats and informing conservation targets

**DOI:** 10.1101/2025.11.01.686047

**Authors:** Manuel Gonzalez-Rivero, Kerryn Crossman, Angus Thompson, Juan C. Ortiz, Sun W. Kim, Michael Emslie, Andrew S. Hoey, Mia Hoogenboom, Kevin Bairos-Novak, Eva McClure, John M. Pandolfi, Peter J. Mumby, Murray Logan, Britta Schaffelke, Timothy Staples, Katharina E. Fabricius

## Abstract

The goal of ecosystem management is to maintain healthy and resilient ecosystems over time. These attributes can be summarised under a general term widely used in management plans: ecosystem condition, defined as the overall quality of an ecosystem relative to a desired or reference state. However, measuring and monitoring ecosystem condition remains a challenge. Monitoring ecosystem condition requires a suite of ecological indicators that simultaneously capture the complexity and variability of ecological processes, while being informative and usable in management and decision-making contexts. Such indicators need to be holistic, measurable, sensitive and scalable. Here we outline a resilience-based monitoring framework for coral reefs that consolidates monitoring data and research insights into a relevant, integrated format. The framework includes indicators of ecosystem state (coral cover) and key processes (represented by recovery performance, macroalgae prevalence, community composition, and coral juvenile density). Indicators are generated and scaled from monitoring data by applying explicit, reef-specific thresholds, providing a simple but comprehensive set of ecosystem condition values. Using cases from the Great Barrier Reef, we demonstrate how the framework integrates these indicators for detailed assessments of reef habitat condition and its potential role in management. Based on these case studies, we discuss important considerations for applying this framework worldwide, acknowledging current limitations related to data availability, resolution, and the length of time series.

## 1 Introduction

Managing the condition of ecosystems is a central tenet of conservation planning, particularly in the context of global change (Chase et al., 2018; Game et al., 2008; Moore and Schindler, 2022; Mumby et al., 2014; Oliver et al., 2015). There are multiple conceptual definitions of condition in ecological systems, but in the broadest sense, condition refers to the ecosystem’s overall quality, measured by whether key attributes are maintained relative to a reference state (Czúcz et al., 2021; Stoddard et al., 2006). From an operational perspective, ecosystem condition is linked to the long-term maintenance of ecosystem structure and function, thereby allowing exploration of terms such as ecological integrity, ecosystem resilience, ecological degradation, and sustainability. In this study, we approach ecosystem condition by considering the capacity of ecological systems to resist change and/or rebuild following stress, key properties that also define ecological resilience (Gunderson, 2000; Oliver et al., 2015).

Operationalising resilience for coral reef management presents three central challenges. First, resilience operates over long temporal scales, wherein ecosystems are dynamic and may transition between different states. Therefore, inferring resilience from conventional monitoring observations requires accounting for multiple potential futures that are uncertain and less predictable (Arani et al., 2021; Mumby et al., 2014), with resilience assessed as the probability that a reef remained in its “desired” state (Arani et al., 2021; Mumby et al., 2014). Second, mechanistically, resilience is underpinned by many processes that contribute to the ecosystem’s response to disturbances, including factors that influence population growth, vulnerability to specific disturbances, and adaptive potential, among others (Oliver et al., 2015). Third, coral reefs are complex hierarchical systems, and changes can manifest across different spatial scales and functional domains (Donovan et al., 2023). Additionally, acknowledging and understanding their fluid and complex nature is of ever-increasing importance to define expectations of a reference state (Moore and Schindler, 2022).

In coral reefs, hard coral cover is a widely used indicator of reef status and trends (Miloslavich et al., 2018) because it is easily measured and understood by a non-technical audience and has been consistently collected worldwide for decades. However, assessing reef condition including its resilience beyond simply describing hard coral cover requires (1) indicators that describe the multiple attributes and processes driving the resistance and recovery capacity of reef ecosystems, (2) an understanding of thresholds that define states across different spatial and temporal scales; (3) recognising the variability of resistance and recovery processes; and (4) a synthesis metric to communicate and evaluate the effectiveness of management strategies. These requirements cannot be captured solely by hard coral cover dynamics. Recent studies have demonstrated that expanding beyond coral cover to define habitat condition improves the quality of conservation actions (Nolan et al., 2021), resulting in transparent spatial prioritisation that meets conservation targets while minimising socio-economic costs (Ball et al., 2021; Moore et al., 2016).

Coral reef communities vary spatially due to biogeography and temporally due to succession dynamics in response to chronic and acute pressures (Brito-Millán et al., 2019; Mumby et al., 2014; Nyström and Folke, 2001; Pisapia and Pratchett, 2014; Tebbett et al., 2022). Eliciting prognostic assessments of future trajectories of coral reef habitats is challenging when evaluating coral cover alone, because multiple processes —such as competition, juvenile replenishment, and taxonomic composition —drive recovery (Nyström et al., 2008). While the resulting coral cover after a disturbance influences recovery dynamics, its effect is mediated by these other processes. Therefore, evaluating coral cover after disturbance can yield contrasting recovery trajectories depending on the status of different indicators. Furthermore, focusing solely on coral cover can mask essential changes to reef composition and diversity (Knowlton, 1992; Mumby et al., 2011). These shifts have been observed worldwide (McWilliam et al., 2020; Pandolfi et al., 2020; Pratchett et al., 2011), potentially affecting ecosystem functions such as carbonate accretion, habitat provision, and recovery rates (Johns et al., 2014a; Perry and Alvarez-Filip, 2019; Richardson et al., 2020). A holistic assessment of reef condition requires (1) a multidimensional assessment of key indicators that evaluate the condition of reef habitat structure (Nyström et al., 2008), (2) evaluating the current state of these indicators against regional or local expectations that acknowledge underlying spatial variability in their condition (Knowlton, 1992; Mumby et al., 2011), (3) recognise the temporal dynamics of coral reefs to identify early warning signs of declining resilience capability.

Building on previous studies of indicators of coral reef condition (Flower et al., 2017; Mumby et al., 2014; Thompson et al., 2020), we outline a quantitative and methodological framework that integrates five indicators of reef condition based on conventional ecological metrics from existing monitoring time series. Collectively, these indicators assess the resilience capacity of coral reef habitats, assist in interpreting monitoring data, assess management targets and communicate the outlook of coral reefs based on current conditions. The study then introduces a quantitative approach to integrate the monitoring data of these five indicators into a classification for habitat condition. Last, we present use cases from the Great Barrier Reef that demonstrate how the approach addresses the challenges of defining and quantifying reef habitat condition and provides prognostic inferences about resilience.

## 2 Methods

### 2.1 The Framework

The framework conceptualised in this study describes a methodology for calculating standardised indicator metrics to evaluate changes in the condition of reef habitats. The methodology follows four steps: (1) identify indicators and metrics, (2) define thresholds for expectations, (3) scale metrics into indicator scores for comparable assessments, and (4) define a logic for the interpretation of integrated index scores using a condition typology (see Electronic Supplementary Material, ESM 1).

Indicators define broad, overarching attributes of resilience that describe the performance of ecological attributes. The framework includes indicators of ecosystem state (coral cover) and key processes (represented by recovery rate, macroalgae prevalence, community composition, and coral juvenile density). To operationalise indicators, metrics were defined to provide specific, quantifiable measurements for assessing their performance (Table 1). A critical element of the indicator framework is the inherent relationship among indicators, which, when combined, provide a holistic assessment of the condition of coral reef habitats. Scaling metrics into scores creates a common reference space for comparing metrics across indicators, which are typically measured in different units and exhibit varying ranges. The methodological approach for this framework is summarised in the sections below, providing references to more detailed methodological approaches in the supplementary materials.

**Table 1.**
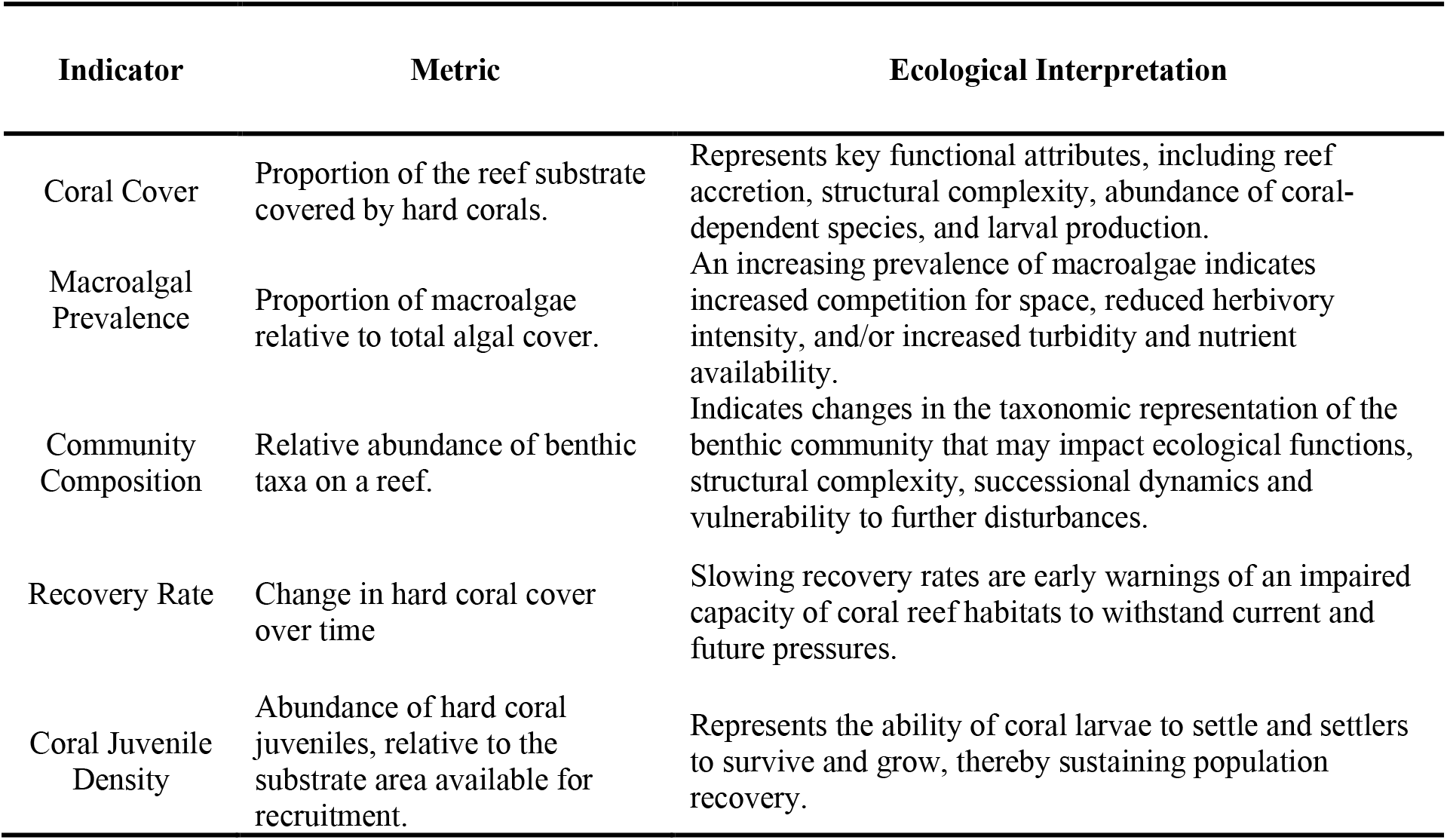
List of indicators included in the Framework with a brief rationale for their inclusion.

| Indicator | Metric | Ecological Interpretation |
| --- | --- | --- |
| Coral Cover | Proportion of the reef substrate covered by hard corals. | Represents key functional attributes, including reef accretion, structural complexity, abundance of coral-dependent species, and larval production. |
| Macroalgal Prevalence | Proportion of macroalgae relative to total algal cover. | An increasing prevalence of macroalgae indicates increased competition for space, reduced herbivory intensity, and/or increased turbidity and nutrient availability. |
| Community Composition | Relative abundance of benthic taxa on a reef. | Indicates changes in the taxonomic representation of the benthic community that may impact ecological functions, structural complexity, successional dynamics and vulnerability to further disturbances. |
| Recovery Rate | Change in hard coral cover over time | Slowing recovery rates are early warnings of an impaired capacity of coral reef habitats to withstand current and future pressures. |
| Coral Juvenile Density | Abundance of hard coral juveniles, relative to the substrate area available for recruitment. | Represents the ability of coral larvae to settle and settlers to survive and grow, thereby sustaining population recovery. |

### 2.2 Indicators and metrics

Five indicators were selected based on stakeholder consultations with managers of the Great Barrier Reef Marine Park, available monitoring datasets and previous research into key mechanistic attributes of resilience (McClanahan et al., 2012; McLeod et al., 2021; Mumby et al., 2013; Thompson et al., 2020). Each indicator was defined by a specific metric (Table 1) calculated from monitoring data, using detailed methodological approaches described in the Electronic Supplementary Materials (ESM 2-6).

### 2.3 Thresholds to define expectations

To interpret ecological indicators commonly measured by monitoring programs worldwide, values must be evaluated against expectations or reference conditions (Flower et al., 2017; Huggett, 2005; Stoddard et al., 2006; Thompson et al., 2020). However, making consistent interpretations of the condition of these habitats at any given time is challenged by (1) the spatial heterogeneity of reef habitats (Levin, 1992; Richards, 2013); (2) data availability to define reference conditions before human settlement (Jackson, 1997; Pandolfi et al., 2003); and (3) a need for understanding the consequences of critical levels of these indicators in terms of the resilience of these systems (Briske et al., 2010; Mumby et al., 2007; Mumby et al., 2011; Osman et al., 2010). Acknowledging these challenges, we established two reference thresholds, maintenance and critical thresholds, to aid in interpreting indicators.

#### 2.3.1 Maintenance thresholds

The maintenance threshold is analogous to a previously defined term, the Minimally Disturbed Condition reference (Stoddard et al., 2006). This threshold identifies the expected value for each indicator using a reference period from coral reef monitoring that is sufficiently long to capture the natural variability of each indicator during a period when ecosystems were less disturbed, compared to recent times. As the term suggests, the maintenance threshold quantifies the desirable state that reef habitats must maintain to function as we know them, based on contemporary observations.

Accounting for the natural spatial variability of reef habitats, the framework used spatially constrained references to calculate a maintenance threshold representative of bioregions. For the Great Barrier Reef (GBR), we used the ‘reef bioregions’ (Fernandes et al., 2005; Kerrigan et al., 2010), defined as 30 discrete environmental provinces representing specific coral reef habitats and biodiversity characteristics. Historical datasets were aggregated by bioregion to estimate the maintenance threshold, except for Macroalgal prevalence, which was estimated at the site level due to high variability within inshore bioregions.

Based on historical observations from monitoring and recorded disturbances in the GBR, we identified the period from 1995 to 2015 as a suitable reference period for estimating maintenance thresholds. This period both preceded an exponential increase in the likelihood of disturbances affecting coral reefs in the GBR (Emslie et al., 2024), while capturing fluctuations in indicator values over time. This reference period was consistent across all indicators and bioregions with monitoring data, except for the Recovery Rate indicator, for which limited recovery-period data for parameterising ecological models necessitated extending the reference period to 2020. Thresholds were calculated as the estimated average distribution of metrics for each indicator using methodological approaches summarised in Table 2 and expanded in the ESM.

**Table 2.**
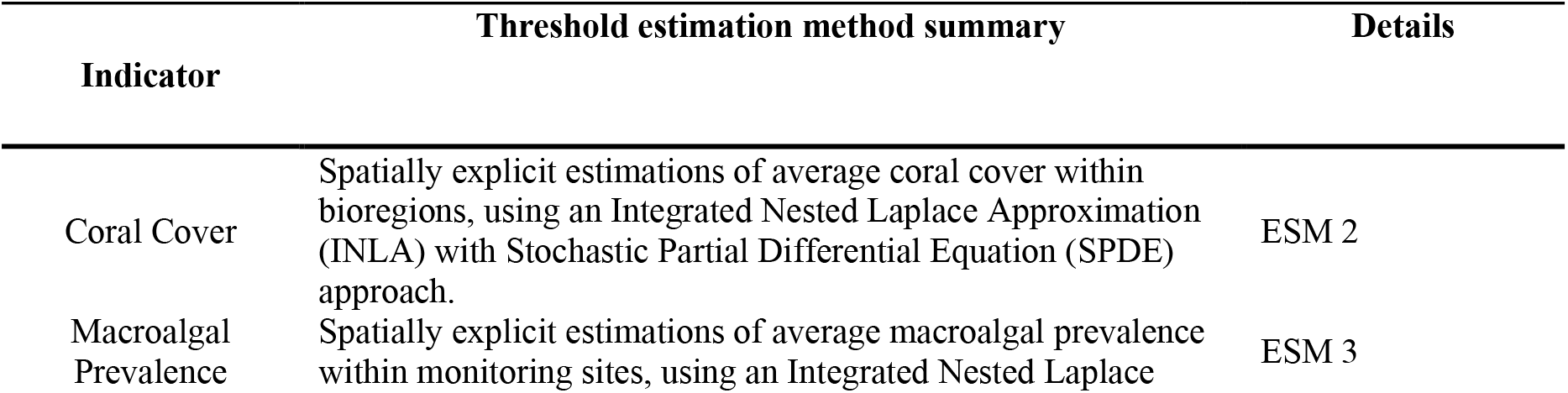
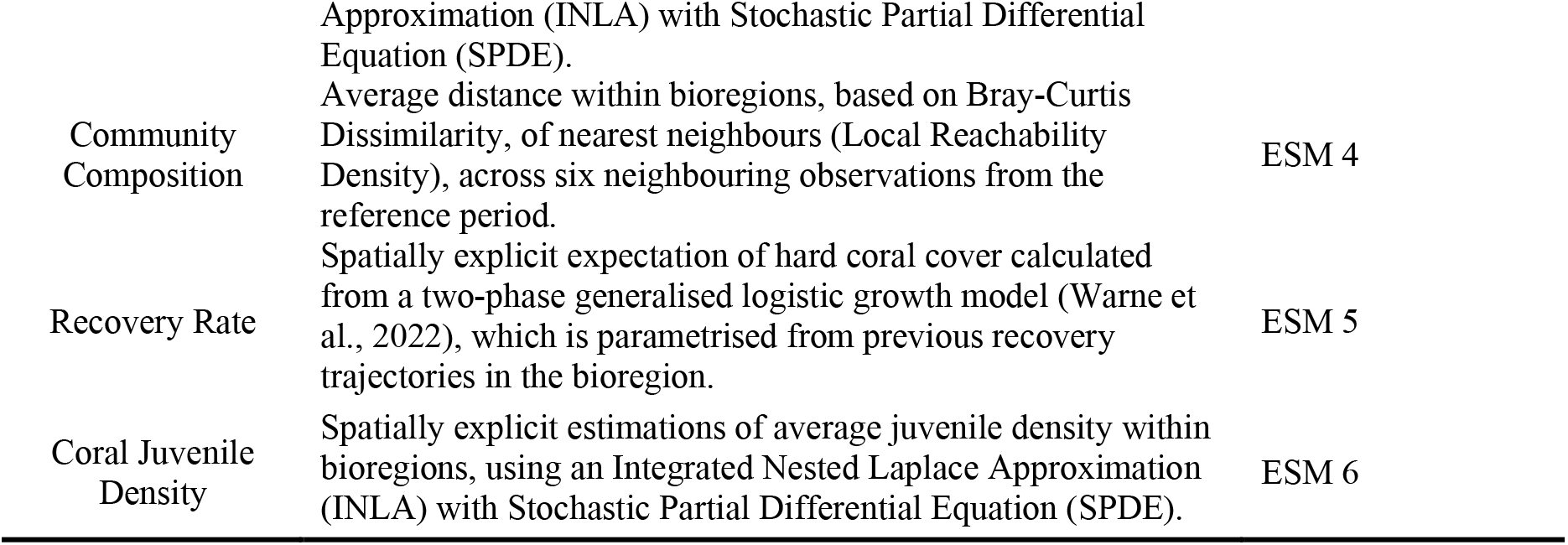
Approach for estimating maintenance thresholds for selected indicators, with details provided in the Electronic Supplementary Materials (ESM)

#### 2.3.2 Critical thresholds

While the maintenance threshold provides a quantitative estimate of expectations, historical observations are temporally constrained to narrow, contemporary periods (e.g., the past few decades), compared with the over 8,000 years of modern reefs’ existence and evolutionary diversification (Roff, 2020). Therefore, reference values from estimated maintenance thresholds may already reflect a shifted baseline or underestimate the natural variability in indicator values across much larger temporal scales (Jackson et al., 2001; Knowlton and Jackson, 2008; Pandolfi, 2002; Pandolfi et al., 2003; Pandolfi et al., 2020).

To address concerns about potentially biased interpretation of indicators based on modern, potentially already perturbed states, a critical threshold was developed to provide a reference below which the ecological attributes represented by an indicator are expected to be severely compromised. The reference values for three of these thresholds have been formulated to determine the minimum expectations for the long-term persistence of key ecological functions (Table 3).

**Table 3.** Reference criteria for critical thresholds of indicators.

| Indicator | Metric | Threshold values | Interpretation |
| --- | --- | --- | --- |
| Coral Cover | Minimum hard coral cover (%) required to ensure positive net carbonate production. | 20% | When hard coral cover is below this threshold, reefs are unlikely to create new habitats, and the reef matrix's existing habitats are likely to erode (i.e., a negative carbonate budget; ESM 2). |
| Coral Juvenile Density | The minimum density of *Acropora* juveniles that is required to maximise the likelihood of recovery. | Inshore:<br>0.54 ind.m<sup>-2</sup><br>Offshore:<br>0.95 ind.m<sup>-2</sup> | Densities of *Acropora* juvenile corals below this value suggest the unviability of key populations supporting the recovery of reef habitats following disturbance (ESM 6). |
| Macroalgae Prevalence | Macroalgae prevalence (see Table 2), beyond which the density of *Acropora* juveniles falls below critical levels. | 22 % | Macroalgal prevalence above this threshold will likely limit the reef's recovery by reducing juvenile abundance (ESM 3). |

Since most coral reef ecosystem functions are founded on the bioconstruction of reefs (Wolfe et al., 2020), identifying whether the reef has a net positive carbonate budget is an intuitive metric to support functionality (Mace et al., 2014). Perry et al (2013) found that an average coral cover of 10% was associated with net carbonate production in Caribbean reef systems. However, rates of fish-based bioerosion are much higher in the Pacific (Mumby and Steneck, 2018), so the required coral cover needed to offset parrotfish bioerosion is higher. There remains considerable uncertainty on the appropriate cover for the Great Barrier Reef, but a recent estimate suggested ∼20% (ESM 2).

Densities of juvenile coral colonies are often used as proxies for recruitment success, as their abundance reflects the cumulative effects of larval supply, settlement, and post-settlement processes, and often correlates with community recovery trajectories (Doropoulos et al., 2015). Among juvenile hard coral taxa, *Acropora* is often proposed as a good and prognostic indicator of coral cover recovery dynamics (Doropoulos et al., 2015; Emslie et al., 2008; Johns et al., 2014a; Roff, 2020). Low densities of *Acropora* juveniles are associated with limited recovery of coral cover and shifts in community composition (Bramanti and Edmunds, 2016; Yadav et al., 2018). However, recognising the spatial and temporal variability of recruitment and population recovery, the minimum density of *Acropora* juveniles required to ensure sustainable population growth and community recovery remains unknown. A recent estimate, using population demographic modelling, suggests that minimal densities of 0.54-0.96 ind.m^-2^ in inshore and offshore habitats are required to ensure coral cover recovery in the GBR (ESM 6).

Macroalgae are ubiquitous in coastal reef habitats and often dominate the benthos of inshore reefs in the GBR (Fabricius et al., 2023). While macroalgal communities often contribute to key ecological functions in coral reefs, values above 20% cover tend to be negatively correlated with coral cover (Ceccarelli et al., 2020), due to their superior competitive outputs and their capacity to pre-empt available space for recruitment (Brown et al., 2020; Jompa and McCook, 2003; Mumby et al., 2005). To provide an assessment of the likely impact on reef recovery dynamics, we use estimates of critical levels of macroalgal prevalence —the relative abundance of macroalgae to total algal cover —at which the juvenile density of Acropora corals is severely constrained (ESM 3).

### 2.4 Scaling metrics into index scores

Index scores were evaluated as the distance from monitoring observations to maintenance and critical thresholds and then scaled to a standard reference space to facilitate comparisons across indicators and geographies. Indicator metrics (e.g., % hard coral cover) are evaluated against their thresholds to calculate the index score, which measures the distance between the observed metric and each threshold. Index scores are scaled from 0-1, enabling consistent interpretation across metrics. Generally, index scores can be interpreted as increasingly good states as they move from 0.5 to 1.0, and increasingly poor states as they decline from 0.5 to 0.

Except for the indicator ‘community composition’ (see ESM3), index scores were calculated using the posterior draws from the indicator model for a given year and reef. Each drawn posterior is paired with a drawn posterior from the maintenance and critical models. The score is calculated for each pair as the difference between the observed value and each reference threshold, then scaled to 0-1 using the cumulative distribution function of a logistic distribution for a given distance metric (see ESM for each indicator). Therefore, when the index score approaches 1, the observed indicator metric is well above its corresponding threshold. Mid-levels (0.5) of the indicator score suggest that the observed indicator metric matches the threshold used to compare it against, and a zero value indicates the highest certainty that the metric is below the corresponding threshold.

### 2.5 Condition typology and interpretation

The framework follows a logical process to estimate index scores for each indicator, identify underperforming indicators, and build an interpretation of the whole-of-ecosystem condition using a typology or classification system (Fig. 1).

**Figure 1.**
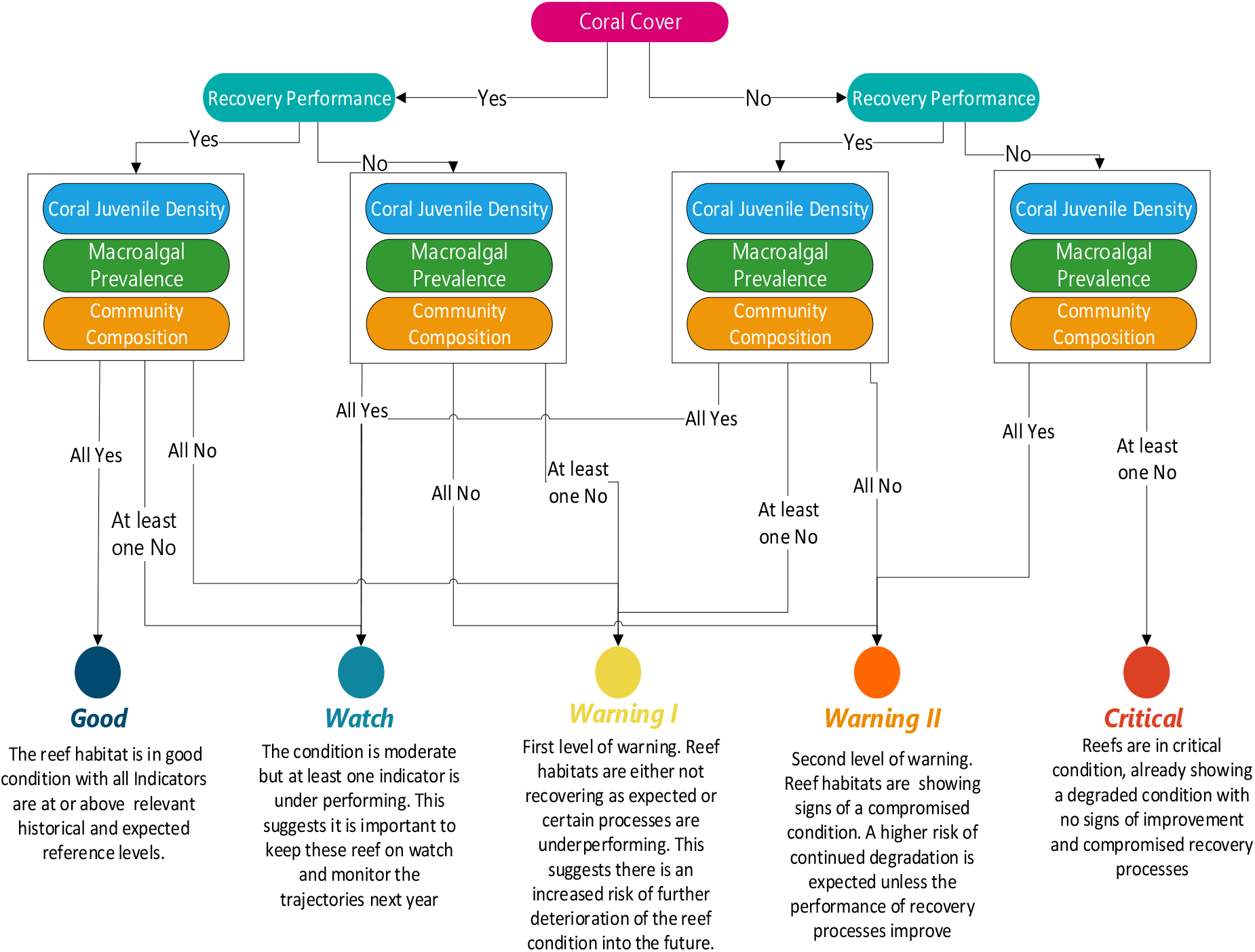
Indicator Framework reef condition classification flow diagram. Yes/no responses concern the following question: Are current indicator scores at or above the maintenance and critical thresholds?

For three of the five indicators (Coral cover, Macroalgae, and Juvenile Coral Density), the critical reference provides an explicit, directional assessment of observed values relative to a threshold below which the processes represented by the indicator are impaired. Consequently, the overall condition is assessed by averaging the two scores for each indicator. To reflect that the critical threshold indicates a severely compromised state of the ecosystem attributes considered here, critical indicator scores below 0.5 are set to 0 before averaging. For the recovery rate and community composition indicators, the critical thresholds are more complex than a straightforward translation to a severely compromised state, so only maintenance scores are used to evaluate reef condition. Across all five indicators, underperforming is detected if the probability that the score is below 0.5 is high (p > 0.8 in this study).

To describe, interpret and communicate the condition of ecosystems from monitoring data, a typology of reef habitat conditions was defined. This typology employs a hierarchical classification that considers each indicator’s role in the ecosystem’s resilience and categorises reef habitat condition into five classes. These classes depict a range of states based on a risk-assessment approach that evaluates the likelihood of ecological underperformance and its potential impacts on resilience. Likelihood is assessed using credible intervals to reflect variability in individual indicator scores. The impact is evaluated via a logical hierarchy of each indicator’s contribution to reef habitat resilience (Fig. 1). If all indicators meet or exceed expected thresholds, the reef habitat is classified as “Good”. However, assessing the ecological impact of other habitat condition classes involves more nuance. When coral cover and recovery rate indicators score below 0.5, reef habitats are considered compromised. Process indicators, such as macroalgal prevalence, juvenile abundance and community composition, enable further categorisation into “Warning I”, “Warning II”, or “Critical”. A “Watch” class was also included to identify situations where only one indicator underperforms, but the overall system is likely to recover. This helps pinpoint reef habitats that need further monitoring to determine whether management actions are necessary.

### 2.6 Framework outputs

When applying the indicator framework at regional scales that aggregate across individual reefs, it is also informative to examine variability in scores among reefs and the number of reefs where changes are evident.

The indicator framework codebase provides a distribution of scores for each metric and reef-year combination in the input monitoring time series. These distributions are summarised to provide medians and the probability that the median score is <0.5, across various scales and condition classes, from individual reefs to aggregations reflecting spatial management units (e.g., GBR Management Zones and Australian Natural Resource Management (NRM) regions) or biogeographical zones (e.g., bioregions and offshore/inshore habitats).

## 3 Results

The described indicator framework enables comprehensive assessments and concise interpretation and reporting of the state and resilience capability of reef habitats, based on routine monitoring data. Below, we provide three worked examples with data from long-term monitoring programs conducted by the Australian Institute of Marine Science. These demonstrate the application of the framework method and illustrate the breadth of applications for assessing and reporting on reef condition at different scales, from individual reefs to management regions, as well as the combined value of the selected indicators for coral reef condition assessments.

### 3.1 Use Case #1: A holistic assessment of habitat condition and biodiversity

#### 3.1.1 Rationale

In addition to measuring and reporting live hard coral cover as the key indicator for coral reef condition, compositional changes over years and decades reveal fundamental changes in reef habitats. The described indicator framework allows for exploring and interpreting changes in coral cover alongside other indicators, identifying potential changes that might affect ecosystem functions such as carbonate accretion, habitat provision, and recovery rates.

#### 3.1.2 Worked example

Agincourt Reef, northern GBR, has been impacted by several disturbances (inc. tropical cyclones, crown-of-thorns starfish outbreaks, and thermal stress) over consecutive years (Fig. 2A). However, in recent years, hard coral cover has recovered to historical maximum levels.

**Figure 2.**
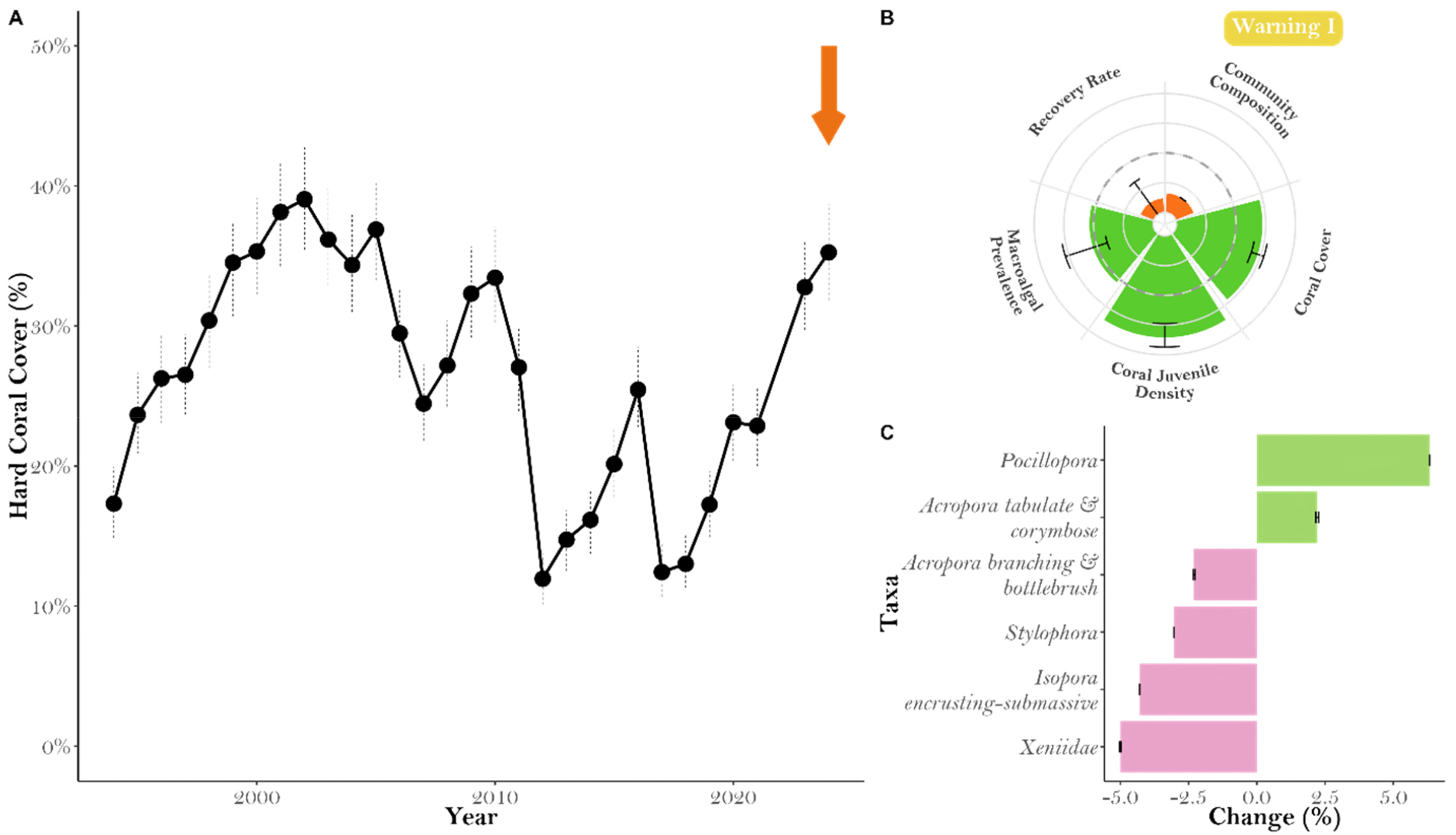
Recovery dynamics of Agincourt Reef (northern GBR), where hard coral cover reached historical maximum levels in 2024 (A, orange arrow), while shifts in community composition and slow recovery rates in 2024 are captured by the framework (B). The radial plot shows the index scores for each indicator. Values below reference levels (0.5, dashed line) are shown in orange, and those within or above the historical averages are shown in green. Major taxonomic changes (gains in green and losses in pink) compared to historical averages allow a deeper understanding of the community changes captured by the indicator (C).

Reviewing this pattern solely based on coral cover masks foundational changes that occurred after these disturbances. The Framework highlights substantial changes in community composition and slower-than-average recovery rates by 2024, while the other indicators remained within expected levels (Fig. 2B). Benthic community changes suggest *Pocillopora* and tabulate and corymbose *Acropora* have increased in abundance compared to historical levels, while other historically abundant taxa have shown substantial losses (Fig. 2C).

### 3.2 Use Case # 2: Prognostic assessment of recovery

#### 3.2.1 Rationale

The frequency and intensity of disturbances profoundly affect the future trajectories of coral reef condition, and prognoses of reef recovery potential are essential for spatial prioritisation in conservation planning. The remaining coral cover after a disturbance is an important predictor of recovery. However, contrasting recovery trajectories are possible, depending on other key processes, such as competition, juvenile replenishment, and taxonomic composition, that influence recovery rates. The indicator framework combines the information from the five indicators to enable differential interpretation of coral cover trajectories for prognostic assessments.

#### 3.2.2 Worked example

Snapper Island, northern GBR, was severely affected by a tropical cyclone in 2014 and a mass coral bleaching event in 2017 (Fig. 3A). By 2018, coral cover remained low. The framework analysis revealed key processes that were underperforming relative to historical baselines, suggesting the reef remained in a compromised state (Fig. 3B). Specifically, macroalgal prevalence was above average, juvenile coral abundance was below average, and community composition had undergone substantial changes. Subsequent monitoring records showed only a marginal recovery of coral cover by 2024.

**Figure 3.**
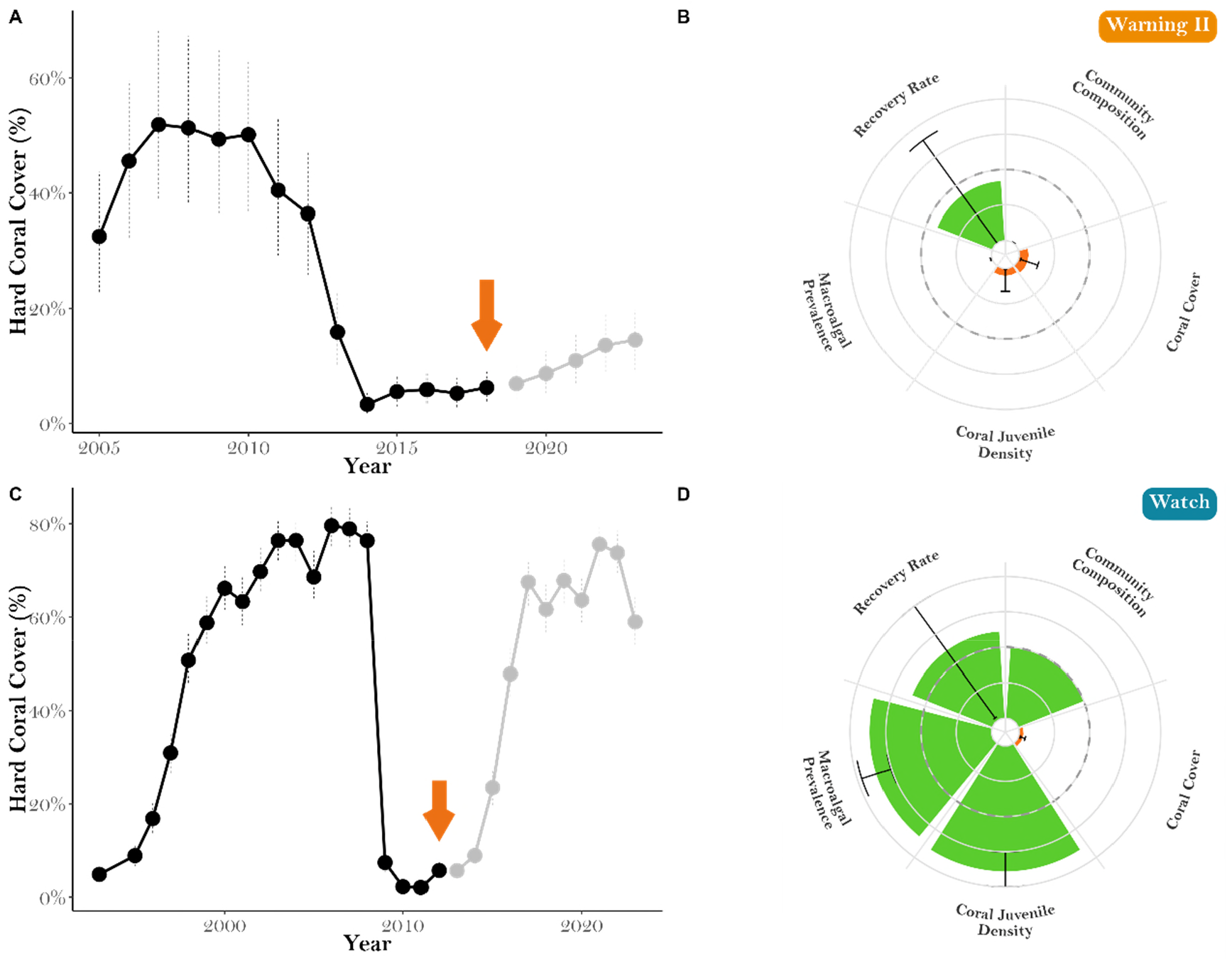
Contrasting examples of coral cover recovery and prognostic assessments from the framework. Temporal trends of coral cover for Snapper Island (A) and Lady Musgrave Reef (C) are evaluated for specific years (orange arrows). The results from applying the Framework for the years indicated by the orange arrows show contrasting interpretations (B and D). Values below historical averages (0.5, dotted line) are shown in orange, those within or above the historical averages are shown in green. The grey lines in the temporal coral cover plots show the observed trajectories after the evaluated year (orange arrows).

3.3 A second location, Lady Musgrave Reef in the southern GBR, was heavily impacted by two severe cyclones in 2008-2010. Evaluating coral cover two years later (2012) shows that it stayed low (Fig. 3C). However, assessing the other four indicators (Fig. 3D) suggests that at that time, the ecosystem maintained key elements supporting its recovery, including juvenile coral abundances, recovery rates, macroalgal prevalence, and community composition, all at or above expectations. Therefore, although hard coral cover was low, the Framework indicates that the reef remained capable of recovery. Fast-forward to data collected in 2024, and it is evident that the reef has shown strong resilience and has regained its historically high levels of coral cover (see more details in the <u>AIMS Monitoring Reef Dashboard</u>).

### 3.4 Use Case #3: Spatial management prioritisation

#### 3.4.1 Rationale

Designation of locations or areas under specific management instruments is central to spatial marine planning and management. This may include marine protected areas, zoning for extractive activities, temporary protection zones, locations for coral reef restoration, and seascape-scale conservation strategies such as managing activities on catchments adjacent to coral reefs or reef fish habitat connectivity from coasts to coral reefs. Information on coral reef habitat condition is often missing in spatial planning, potentially leading to the prioritisation of actions in less threatened or lower-quality habitats. The indicator framework expands beyond coral cover, providing relevant information about coral reef habitat condition for spatial prioritisation applications.

#### 3.4.2 Worked example

In the Great Barrier Reef, the Reef 2050 Long-Term Sustainability 2021-25 Plan serves as an overarching instrument that sets the direction of conservation strategies and management actions in the GBR World Heritage Area (GBRWHA). Key objectives of the Plan include maintaining coral reef habitats in good condition and maintaining their resilience. Based on this premise, this case study illustrates the application of the indicator framework to assess the condition of reef habitats across NRM regions, habitats and indicators.

Condition scores for coral reef habitats in 2024 across the GBR were aggregated for each of the six NRM regions within the GBRWHA, additionally divided into inshore and offshore habitats (Fig. 4A). The analysis showed that inshore coral reef habitats in most NRM regions in 2024 were in a compromised condition between Warning I and Critical, except for the Wet Tropics NRM. For example, using data from monitored reefs in the Fitzroy NRM region allows managers to identify the reefs (Fig. 4B) and the indicators (Fig. 4C) that most significantly contribute to the regional classification of Critical. Overall, the results aligned with previously reported analysis of inshore monitoring data from coral reef habitats and water quality parameters. According to these reports, coral reefs in the Fitzroy NRM region have shown a declining trend, with the lowest reef condition estimate over the past 20 years, a result of a combination of recent marine heatwaves and likely chronic effects from specific water quality parameters (Thompson et al., 2025). Recent water quality analyses show high concentrations of total suspended solids and excess nutrients, in particular, oxidative forms of nitrogen (Moran et al., 2025), which also corresponds to accelerated rates of land clearance in this region (Venegas-Li et al., 2025). The analysis using the indicator framework supports the identification of priority regions to guide resource investment for various conservation efforts.

**Figure 4.**
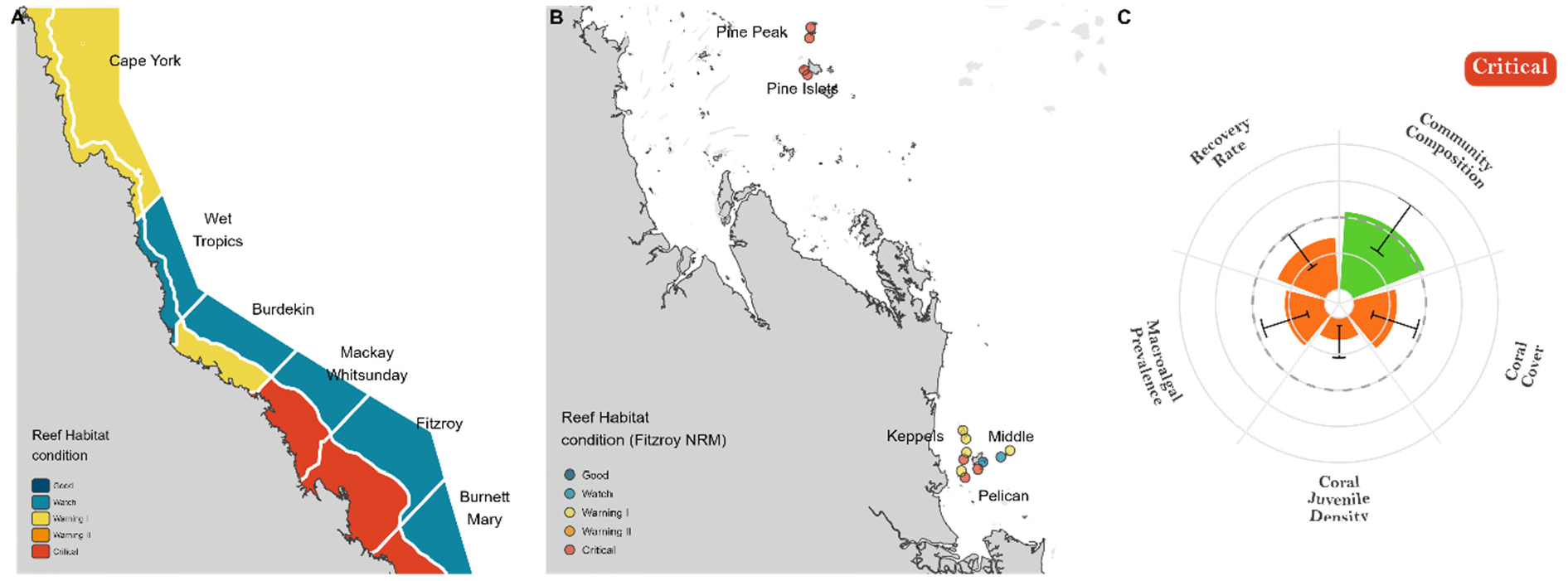
Spatially aggregated classification of reef habitat condition in 2024 for each Natural Resource Management (NRM) region in the Great Barrier Reef and Inshore and Offshore habitats (A). Reef-level assessment across the Inshore Fitzroy RMN (B) and across indicators (C) allows a deeper evaluation for management prioritisation.

## 4 Discussion

A recurrent challenge to improving the efficiency of ecosystem management in executing effective actions is access to actionable information that triggers decisions and drives policymaking (Cook, 2016; Svancara et al., 2005). Effective ecosystem-based management requires deciding which attributes reflect ecosystem-state changes, synthesising them into higher-level metrics, and contextualising these metrics relative to desired ecosystem states and functions (Hennessey et al., 2025; Johnson, 2013; Levin and Möllmann, 2015). Our quantitative framework and a coral reef condition classification support the interpretation of monitoring data and guide the delivery of information for management prioritisation and evaluation. The framework describes coral reef habitat condition by systematically evaluating key indicators that contribute to coral reef resilience, an approach analogous to diagnosing health through symptomatology. This study demonstrates the application of the condition framework in guiding the delivery of actionable scientific information for management through three use cases. The examples provided in this study illustrate a more systematic, holistic, and prognostic evaluation of ecosystem condition beyond assessments based solely on coral cover.

### Novel methodological contributions

The study contributes to expanding the application of ecological threshold concepts for management (Huggett, 2005) and methodological improvements for assessing reef habitat condition (Flower et al., 2017; Thompson et al., 2020). This work builds on the Coral Index, a quantitative methodology for evaluating the condition of inshore and coastal coral reef habitats (Thompson et al., 2020). The framework presented in this study expands the target habitats considered to include inshore and offshore reefs, better represents spatial variability, incorporates proxy assessments of critical ecological functions, and draws on other approaches to assess the contribution of indicators to reef habitat condition.

Building on the Coral Index, the framework considers reference-state thresholds for each indicator across bioregional provinces to provide greater granularity in assessing ecological condition, while recognising the scale-dependent variability of ecosystem thresholds (sensu Spake et al., 2022). Two thresholds are considered in the proposed framework. A first threshold is calculated based on historical values of indicators, prior to a documented increase in the magnitude, frequency, and extent of environmental pressures on coral reefs (Emslie et al., 2024), to define expectations for maintaining the condition of reef habitats as known from modern observations. A second threshold allows for the discretisation of critical indicator levels. It guides the interpretation of early warning signals when a system is likely to undergo abrupt shifts from its current state or to lose resilience potential (e.g., by slowing recovery capacity).

Additional ecological dimensions are included to assist in the interpretations of indicators (e.g., critical thresholds for *Coral cover* and *Juvenile density*) by capturing attributes of ecosystem functioning, such as carbonate accretion and erosion (Bellwood et al., 2019; Perry and Alvarez-Filip, 2019), and the population viability of *Acropora* as a keystone taxa (Ortiz et al., 2021; Wilson et al., 2019). In addition to the added dimensions in evaluating reef condition, the proposed framework also refines and expands previous approaches to improve the assessment of recovery rate (MacNeil et al., 2019; Thompson et al., 2020), to account for non-linear behaviours of coral colony growth rate leading to multiphasic recovery phases in coral reefs (Warne et al., 2022).

### Implementation of the proposed framework to deliver actionable insights for management

Evidence-informed decision-making can help catalyse the development and implementation of effective actions (Salafsky et al., 2019; Walsh et al., 2015). However, embedding evidence into decision-making is challenged by limited time availability from decision-makers to synthesise sources of information, time pressures for agile decision-making, access to data and information, understanding the Value of Information, and access to information in an actionable form (Bolam et al., 2019; Christie et al., 2020; Pullin et al., 2020). Through use cases, the framework illustrates the utility of spatially explicit and prognostic assessments of coral reef condition, facilitating the prioritisation of reef habitats and management regions for conservation while easing communication of the complex responses of these ecosystems to disturbances. By integrating different lines of ecological “evidence” from coral reef monitoring into a knowledge translation framework, the resulting evaluation of condition improves the quality, actionability, and readiness of available information (sensu Kadykalo et al., 2021; Pullin and Knight, 2003).

Coral reef management faces a myriad of decision-making challenges that vary across management objectives and targets, spatial scales and timeframes (Anthony et al., 2015; Comte and Pendleton, 2018). In the GBR Marine Park, for example, there are several opportunities to implement the proposed framework to enhance current procedures for spatial prioritisation of management strategies and to evaluate the effectiveness of these actions. For example, evidence suggests that prioritising culling efforts to control Crown-of-Thorns starfish outbreaks based on the ecological value and condition of reefs can have profound impacts on the efficacy of managing these outbreaks (Castro-Sanguino et al. 2023, Matthews et al. 2024) and on the framework to complement existing site and regional prioritisation processes. In this case, ecological value is defined as the potential to provide coral larval supply to reefs that have been heavily damaged and in need of replenishment (Mumby et al., 2021).

Managing the impacts of land use on coastal ecosystems is complex and multidisciplinary, spanning social, economic, and ecological perspectives (Halpern et al., 2008; Sahavacharin et al., 2022). Acknowledging that ecological prioritisation is one attribute to consider in formulating complex management decisions, the framework supports the delivery of compelling, high-level and outcome-centred insights from monitoring to inform the prioritisation and evaluation of land-sea management efforts that account for uncertainty (sensu Brown et al., 2019). Further to land-use, climate change is increasingly demonstrated to interact with local pressures(Harvey et al., 2018). As shown in Use Case # 3, the framework allows evaluating the outcomes from cumulative pressures as evidence to prioritise multiple interventions under an adaptive management response to environmental disasters (sensu Harvey et al., 2018; Streit et al., 2025). Lastly, alongside spatial prioritisation, the framework can contribute to a quantitative assessment of management progress towards desired conservation outcomes, improving transparency, governance efficiency, and the adaptability of overarching conservation instruments, such as the Reef 2050 Plan (sensu Vella et al., 2024).

### Limitations and further improvement

Increased turnover rates in marine communities have been associated with rapid increases in environmental pressures that are restructuring biodiversity (Cunningham et al., 2025; Dornelas et al., 2014; Edmunds et al., 2014). However, changes in structure do not necessarily imply changes in function. While functional compositional changes are also deemed taxonomically novel (Cunningham et al. 2025), functional stability can persist despite taxonomic turnover through species redundancy (Biggs et al., 2020; Purschke et al., 2013). Therefore, the consequence of compositional shifts can have either positive or negative impacts on biodiversity (Chase et al., 2020) and ecosystem functions (Edie et al., 2018). Importantly, for evaluating reef habitat condition, the community composition indicator in its current format does not allow interpretation of directionality from an observed change (e.g., Good or Bad). Instead, it flags taxonomic shifts for further investigation by inferring from structural changes (Use Case # 1). Using functional traits (sensu McWilliam et al., 2020) or taxonomic richness (sensu Edie et al., 2018) to estimate reference values for a critical threshold will improve the current framework and assist in interpreting the consequences of emerging taxonomic configurations for describing the condition of coral reef habitats.

Following a major disturbance, ecological processes are often influenced for years after the event, which can confound the interpretation of indicator scores. While the framework can provide prognostic insights into recovery from the condition, it is essential to evaluate reef condition over multiple years post-disturbance to assess reef trajectories. For example, recovery rates cannot be estimated for the first observation following a major disturbance, as there is no logical expectation of an increase in coral cover relative to historical or contextual references. In this situation, the index score will relate to the previous disturbance period. Similarly, abrupt changes in macroalgae can occur due to disturbance, as algae are either themselves impacted (e.g., cyclone mechanical removal of algae) or, conversely, bloom due to available space and excess nutrients after the disturbance (Doropoulos et al., 2014). Lastly, following a major disturbance, communities are expected to change in composition. Succession and reassembly of these communities can take multiple years (Johns et al., 2014b); therefore, it is essential to understand the long-lasting effects of biodiversity restructuring (Cunningham et al., 2025).

### Considerations for a global implementation of the proposed framework

Conceptually, the framework lays the foundation for a quantitative, standardised assessment of reef condition, using a typology comparable worldwide and broadly following recommended criteria (Czúcz et al., 2021; Streit et al., 2025). However, implementing this framework to assess reef condition must acknowledge the need for long-term data availability across indicators and methodological consistency across monitoring programs. As these requirements are not always available worldwide (Souter et al., 2021), the framework implementation will need to be adapted to the readiness of monitoring and management expectations.

Coral reef habitats in the GBR have been consistently monitored for two to four decades using methodologies that are comparable, quality-controlled, and assured across monitoring programs. The indicator framework was designed to capitalise on the availability of this long-term, high-quality monitoring dataset. Furthermore, detecting early trajectories of coral change after disturbance in the recovery rate indicator relies on a high level of repeatability in coral cover assessments, which is achieved using permanent transects in the selected GBR monitoring programs.

While the robustness, completeness, and longevity of monitoring in the GBR set an ideal standard, it should not deter practitioners from tailoring the proposed framework to available data and knowledge. For example, long-term averages of Macroalgae and Hard Coral cover for large areas (such as Marine Ecoregions of the World) can help establish thresholds that can be adjusted based on mechanistic assessments supported by published research or expert elicitation (sensu this work and Gudka et al., 2024). Although inconsistent and detailed taxonomic assessments limit the ability to estimate an indicator score for community composition, data from other indicators can still offer valuable insights into reef condition beyond coral cover alone. Therefore, modifying the framework to operationalise reef habitat condition assessments using readily available information can serve as a practical instrument for guiding management while emphasising the importance of documenting other metrics. Ultimately, while the study demonstrates the value of standardised collection of detailed monitoring data to guide conservation actions, a globally effective use of science to inform management requires higher-quality, more in-depth information generated from global monitoring efforts.

## Supporting information

ESM 1

ESM 2

ESM 3

ESM 4

ESM 5

ESM 6

## 5 Acknowledgements

The authors are grateful for the support from the Great Barrier Reef Marine Park Authority in guiding the design of the framework for management applications. This work was supported by the partnership between the Australian Government’s Reef Trust and the Great Barrier Reef Foundation, with support from the Australian Institute of Marine Science, the University of Queensland and James Cook University.

## Author Contributions

MGR led the development of the manuscript. MGR, KC, AT, JCO, SK, ME, ASH, MH, KBN, EM, JMP, PJM, ML, BS, TS and KEF contributed to the Conceptualisation, Formal Analysis, Investigation, and Methodology. KC, AT, KBN, EM and MH contributed to the data curation. MGR, KC, AT, SK, PJM, ML and TS designed and edited the framework codebase. All authors contributed substantially to the study design, manuscript writing and revisions.

## Data and code availability

The datasets used to estimate thresholds and produce use cases in this manuscript are available from the <u>AIMS metadata catalogue</u> (AIMS, 2023), including the <u>Long Term Monitoring Program</u>, the <u>Marine Monitoring Program</u>, the <u>Southern Inshore Zone Monitoring Program</u>, co-funded by the Mackay Whitsunday Isaac Healthy Rivers to Reef Partnership, and <u>ancillary research data</u> used for contextual thresholds. The methodological framework defined in this study was developed in R, and the source code for generating indicator scores from monitoring data is available on <u>GitHub</u>

