## Supplementary material for "Beyond Coral Cover A framework for assessing the condition of coral reef habitats and informing conservation targets": ESM 1

ESM # 1: Methodological overview

### 1 Data Collection

Data for all models are from fixed-site surveys conducted as part of the AIMS Long-Term Monitoring Program (AIMS 2017) and the Marine Monitoring Program (AIMS 2014). Additional inshore monitoring data contributed to establishing maintenance thresholds for the Coral Cover Indicator (Ceccarelli et al. 2020). For detailed descriptions, please see the references for each monitoring program and the [AIMS standard operating procedures](https://www.aims.gov.au/research-topics/monitoring-and-discovery/monitoring-great-barrier-reef/reef-monitoring-sampling-methods). Complementary research datasets used for critical thresholds are detailed in the Electronic Supplementary Materials (ESM 2-6) for each indicator and the data record for this study (AIMS 2023).

### 2 Index Score Calculation

The methodological descriptions to calculate index scores for each indicator generally follow these steps:

1) Establish spatially discrete *maintenance thresholds*
2) Establish a *critical threshold*
3) *Model temporal observations* of the indicator at reef-level
4) Calculate the *maintenance threshold index score*
   a) Set minimum and maximum values (caps) of the indicator that equate to scores of 0 and 1.
   b) For posterior distributions of temporal observations and thresholds, calculate the distances between the temporal observations and the threshold using a Cumulative Distribution Function. Rdscale the distance between the minimum and maximum caps, setting a score of 0.5 to be equal to the threshold.
5) Calculate the *critical threshold index score*:
   a) Set a value of the indicator that equates to a score of 1.
   b) For posterior distributions of temporal observations and thresholds, calculate the distance between the temporal observations and the threshold using a Cumulative Distribution Function. For distances above 0.5, rescale the distances between the threshold and the cap to scores between 0.5 and 1.
   c) Convert distances below 0.5 to an index score of 0.

For the Recovery Performance Indicator, only steps 1) to 4) apply. Calculating the index score for the Community Composition Indicator, however, follows a unique process, since it is based on multivariate data. See the individual Electronic Supplementary Materials for a detailed methodology for each indicator.

### 3 Scores for condition classification

For the Coral Cover, Macroalgae Prevalence, and Coral Juvenile Density indicators, a combined score is calculated at the posterior level as the average of the maintenance and critical scores.

### 4 Aggregation across spatial scales

Maintenance (Recovery Rate and Community composition indicators) and Combined scores (Coral Cover, Macroalgae Prevalence and Coral Juvenile Density indicators) are established at broader spatial scales by aggregating the posterior distributions of index scores across the corresponding reef/habitat models for a given year.

### 5 Thresholds for condition classification

Reef or region condition classification based on condition typology uses credible intervals of scores as thresholds. For the Coral Cover, Macroalgae and Juvenile Density indicators, the credible intervals of the Combined Score are used in the condition classification assessment, while for the Recovery Performance and Community Composition indicators, the credible intervals of the maintenance score are used. Credible intervals can be selected based on the user’s preferences. To aid this process, for each posterior distribution at each aggregation level, we provide the probability that the median of the score is < 0.5. In our case study, we classified indicators as underperforming if the probability of the index score being below 0.5 was above 0.8.
