## Supplementary material for "Beyond Coral Cover A framework for assessing the condition of coral reef habitats and informing conservation targets": ESM 2

ESM # 2: Coral Cover Indicator

### 1 Estimation of thresholds

#### 1.1 Maintenance thresholds

Maintenance thresholds were estimated as the average of historical observations (1995-2015) for each reef bioregion in the GBR (Fernandes et al. 2005, Kerrigan et al. 2010) and categorical survey depth (“shallow” and “deep”). For bioregions with small sample sizes across the monitoring datasets, they were grouped based on geomorphological, environmental, and ecological descriptions when appropriate.

We applied the Integrated Nested Laplace Approximation (INLA) framework combined with the Stochastic Partial Differential Equation (SPDE) approach to estimate proportional hard coral cover. Coral cover was modelled from proportional observations (*p_i_*) using a Binomial Generalised Linear Model (GLM) with a spatially structured random effect ($\varphi_{s_{i}}$), that captures spatial autocorrelation. This component was approximated using a Gaussian Random Field (GRF) through the SPDE approach. To account for variability introduced by differences in survey methodology, temporal fluctuations, and site-specific effects, we included additional random effects for dataset, year, and site. These components help isolate spatial patterns from other sources of variation (see Eq. 1).

${logit}_{p_{ijt}}\sim\beta_{0}+ \varphi_{s_{i}}+ \upsilon_{i}+ \upsilon_{t}+ \upsilon_{j}+\varepsilon_{ijkt}$	(equation 1)

Where:

- *P_ijt_* is the proportion of hard coral cover,
- $\beta_{0}$ is the intercept,
- $\varphi_{s_{i}}$ is the spatially structured random effect estimated through SPDE,
- $\upsilon_{i}$is an i.i.d. random effect for monitoring datasets,
- $\upsilon_{t}$ is an i.i.d. random effect for years,
- $\upsilon_{j}$ is an i.i.d. random effect for reef/habitat
- $\varepsilon_{ijt}$ is the observation-level random effect.

Sites were the unit of sampling for the Gaussian random field. The spatial domain was discretised into a lattice mesh using Delaunay triangulation, with nodes defined by the observed locations, and the boundary was constrained using a shapefile of the GBR. A minimum node distance for the triangles was set to ensure that sites within reefs were in the same triangle (cut-off of 0.95/20). The mesh was then used to define the SPDE.

To establish maintenance thresholds for coral cover at a bioregional level (per depth), a hexagon grid was overlaid onto the sample area (hexagon width approximately 3km, an area approximately 564 hectares). Hexagon areas that sit over a reef were filtered, and the reef area within each hexagon was calculated. Approximate posterior predictions were generated for each hexagon from 1,000 posterior draws. Posterior predictions at the bioregion level represented a weighted (proportional reef area) mean across the hexagons.

#### 1.2 Critical thresholds

The critical threshold for coral cover reflects the minimum coral cover required for a positive carbonate budget, calculated using a carbonate budget model from the [R package “caRbs”](https://github.com/aadesbiens/caRbs).

The model estimates four distinct processes contributing to the net carbonate balance of coral reefs in the Great Barrier Reef (GBR): primary production, secondary production, primary erosion, and secondary erosion.

To estimate carbonate budgets across AIMS LTMP locations, average site-level benthic cover and fish density data from 2016 to 2022 were used as input to the caRbs functions. These data capture a time frame during which size data were collected during visual census surveys of reef fish, an essential criterion for calculating bioerosion through parrotfish feeding. The data also do not include inshore reefs, where visual census surveys of reef fish are not conducted due to visibility limitations.

Site-level primary production was estimated by coral growth, calculated as a function of the mean per cent cover of 56 coral taxa. Taxonomic groups comprised 50 genera, with the genus Acropora further divided into 7 distinct growth forms. Secondary production was estimated by combining per cent cover data for crustose coralline algae (CCA), calcareous red macroalgae, green macroalgae and *Peyssonnelia* with their respective calcification rates, compiled from the published literature when available (Mccormack 2014, Castro-Sanguino et al. 2020, Kennedy & Diaz-Pullido, unpublished data)

Primary bioerosion was calculated based on densities of 23 parrotfish species. To avoid overestimating the contribution of the bump-head parrotfish, *Bolbometopon muricatum*, to bioerosion (due to their large home ranges), its abundance was rescaled from that averaged at a transect scale (250 m2) to its home range of 1 km2. Secondary erosion rates of micro- and macroborers were established from published work that tracked changes in the density and volume of carbonate blocks across locations in the GBR (Osorno et al. 2005, Tribollet and Golubic 2005). Rates of secondary bioerosion within coral skeletons were taken from the literature and stratified across the GBR, whereas rates of surface sponge erosion were linked to observed sponge cover.

A measure of substrate rugosity was obtained by converting ‘substrate complexity’ estimates from the LTMP data, from an ordinal scale of 1-5 to the linear distance covered by a 10 m length of chain fitted to the reef contour (Wilson et al. 2007).

To account for uncertainty, a Monte Carlo simulation approach was used. Within each function, production and erosion rates are randomly drawn from Normal distributions defined by the mean and SD of fitted relationships, derived either by simulation or from published literature, across 10,000 simulations. The resulting outputs are defined as posterior probability distributions (median ± 95% credible intervals). The distributions of each distinct function are then summed to obtain an overall estimate of the carbonate budget for each location.

There was variability in the relationship between coral cover and the carbonate budget among bioregions (Figure S1.1). However, the drivers of the differences are likely complex and levelled out at broader spatial scales. Therefore, a GBR-wide threshold was selected as a sensible approach. At a GBR-wide scale, with no sediment reincorporation to the reef, average coral cover needed to reach 27% for the majority of reefs (80%) to have a positive carbonate budget (Figure S1.2a). However, following the standard approach of assuming that 25% of parrotfish-derived sediment and 50% of the outputs of macro-bioerosion are not lost from the reef, the critical threshold drops to 17% (Figure S1.2b). Given that sediment reincorporation is the global standard but its estimation is highly uncertain, a threshold of 20% was selected to conservatively estimate the minimum hard coral cover required for a positive carbonate budget.

Ongoing work to address the limitations highlighted here, among others, and to improve the modelling approach is expected to result in a revision of this threshold.

### 2 *Estimations of* observation metrics

We used a Bayesian hierarchical linear model to estimate hard coral cover for each observation across years and reef-depth combination (shallow and deep). The response variable was modelled using a beta-binomial distribution to account for overdispersion in coral cover proportions (pi), with a binomial distribution applied to the residuals. Both components used a logit link function and included hierarchical random effects ($\upsilon$) of *reef-depth, site* and *transect* to capture unstructured variability. Including the report year (*year*) as a population-level effect ensures consistency in hindcasting as new years are added to the model, improving temporal comparability and predictive robustness (eq. 2).

${logit(p}_{ijkt})\sim\beta_{0}+ \beta_{1}year+ \upsilon_{k}+\upsilon_{jk}+\upsilon_{ijk}+ \varepsilon_{ijkt}$	(equation 2)

Where:

- *p_ijkt_* is the proportion of hard coral cover for transect *i*, at site *j*, within reef/habitat *k*, in year *t*,
- $\beta_{0}$ is the intercept,
- $\beta_{1}year$is the population-level (fixed) effect of report year,
- $\upsilon_{k}$ is an independent and identically distributed (i.i.d.) random effect for reef-habitat,
- $\upsilon_{jk}$is an i.i.d. random effect for site, nested within reef/habitat,
- $\upsilon_{ijk}$ is an i.i.d. random effect for transect, nested within site,
- $\varepsilon_{ijkt}$ is the observation-level random effect.

For each temporal reef/habitat model, estimates of hard coral cover in each survey year were derived as 1000 draws from the posterior distribution.

### 3 Index Score Calculations

#### 3.1 Maintenance index score

Index scores were calculated using the posterior draws $\left( I_{(i,j)} \right)$ from the indicator model for a given year ($i$) and reef-habitat ($j$). Each drawn posterior is paired with a drawn posterior from the maintenance model ($T_{m}$). The maintenance index score ($f(x)$) is calculated for each pair from the difference between observed values and reference threshold, distance (*x*; Eq. 4), and scaled from 0-1 using a Cumulative Distribution Function of a logistic distribution for a given distance metric (Eq. 3).

$f\left( x;0,1 \right)= \frac{1}{\left( 1+ e^{\left( -x \right)} \right)}$	(equation 3)

Where:

$x= {log}_{2}\left( \frac{I_{(i,j)}}{T_{m}} \right)$	(equation 4)

Minimum hard coral cover value of 0% and a maximum value of 80% were set to index scores of 0 and 1, respectively. Using this approach, a hard coral cover value equal to the maintenance threshold yielded a score of 0.5. Scores above 0.5 were rescaled to extend the upper value of the scores to 1 when the temporal reef value of coral cover was 80%. Scores below 0.5 were not rescaled, as scores naturally are equal to 0 when coral cover is 0%.

#### 3.2 Critical Index score

Index scores were calculated using the posterior draws $\left( I_{(i,j)} \right)$ from the indicator model for a given year ($i$) and reef-habitat ($j$). Each drawn posterior is paired with a drawn posterior from the critical threshold ($T_{c}$). The critical index score ($f(y)$) is calculated for each pair from the difference between observed values and reference threshold, distance (*y*; Eq. 6), and scaled from 0-1 using a Cumulative Distribution Function of a logistic distribution for a given distance metric (Eq. 5).

$f\left( y;0,1 \right)= \frac{1}{\left( 1+ e^{\left( -y \right)} \right)}$	(equation 5)

Where:

$y= {log}_{2}\left( \frac{I_{(i,j)}}{T_{c}} \right)$	(equation 6)

Consistent with the maintenance index approach, the maximum hard coral cover value that equated to a score of 1 was set to 80%, and the critical hard coral cover threshold established from the model described earlier (20%) was set to a score of 0.5. A hard coral cover value equal to the critical threshold yields a score of 0.5. Scores above 0.5 were rescaled to extend the upper value of the scores to 1 when the temporal reef value of coral cover was 80%. In contrast to the maintenance calculation, distances below 0.5 were set to 0 in the critical score calculation.

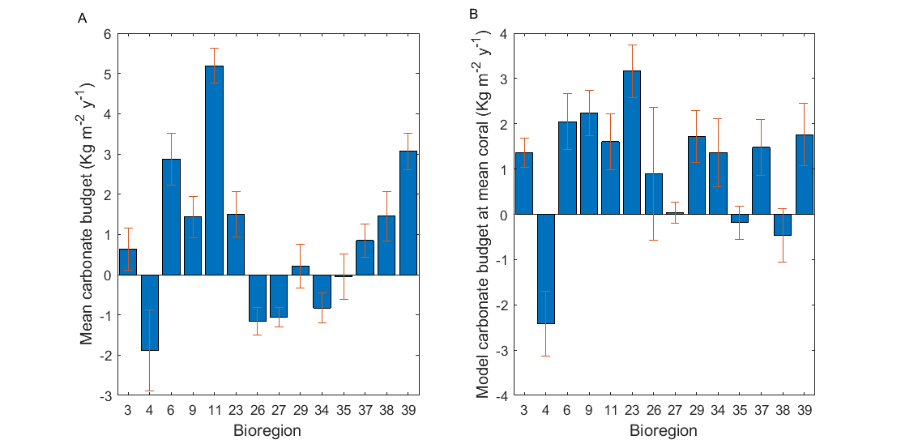

**Figure S1.1:** Differences in carbonate budgets among bioregions of the GBR from 2016-2022. Mean carbonate budget per bioregion (A) and bioregional differences in the underlying relationship between coral cover and budget (B). Error bars denote SEM.

| **A** | 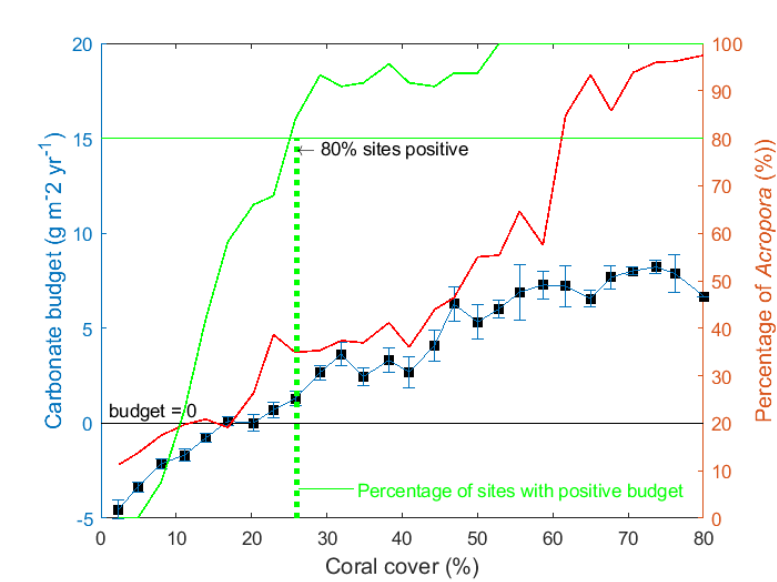 |
| --- | --- |
| **B** | 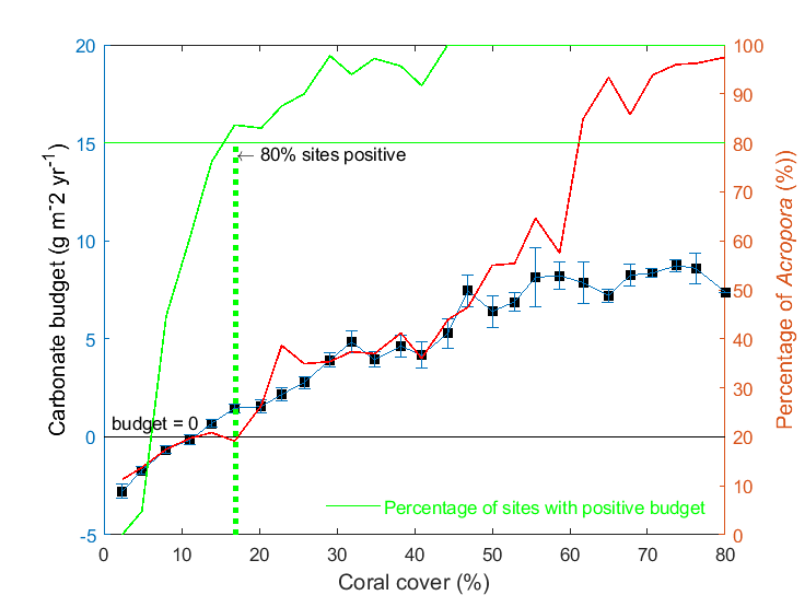 |
| **Figure S1.2:** Threshold level of coral cover needed to achieve a positive carbonate budget (dotted green line) (A) without allowing and (B) allowing for standard reincorporation of sediment into the reef framework. | |
