## Supplementary material for "Beyond Coral Cover A framework for assessing the condition of coral reef habitats and informing conservation targets": ESM 3

ESM # 3: Macroalgal Prevalence Indicator

### 1 Estimation of thresholds

#### 1.1 Maintenance thresholds

For the macroalgae prevalence indicator, the maintenance metric (the proportion of macroalgae within total algal cover) evaluates whether the relative representation of macroalgal species in coral reef algal communities (MAp) is unusual compared to predicted levels based on historical data. Since macroalgae tend to be more prevalent on reefs in turbid, nearshore waters, MAp maintenance thresholds were set at the reef level to accommodate the variability observed in MAp at this finer spatial scale.

Maintenance thresholds of MAp were predicted from spatial models separately for shallow and deep habitats on inshore reefs and for deep habitats on offshore reefs using data from 1995 to 2015. An INLA–SPDE (Stochastic Partial Differential Equation) method was used to model the response of MAp for transects (the sampling unit) and fit it to a binomial distribution. The historical reference model comprised an intercept and a Gaussian spatial field ($\varphi_{s_{i}}$), approximated by a covariance kernel and a Gaussian Markov Random Field (GMRF), which is itself approximated using SPDE. To account for variability introduced by differences in survey methodology, temporal fluctuations, and site-specific effects, we included additional random effects for dataset, year, and hierarchical effects from reef, site and transect (eq 1). These components help isolate spatial patterns from other sources of variation

${logit(MAp}_{ikjt}) \sim\beta_{0}+ \varphi_{s_{i}}+ \upsilon_{ik}+ \upsilon_{t}+ \upsilon_{ikj}+ \varepsilon_{ijkt}$ (equation 1)

Where:

- $\beta_{0}$ is the intercept
- $\varphi_{s_{i}}$ is the spatially structured random effect estimated through SPDE,
- $\upsilon_{ik}$ is the independent and identically distributed (i.i.d.) random effect for site nested within reef-depth
- $\upsilon_{t}$ is the i.i.d. random effect for year
- $\upsilon_{ijk}$ is the i.i.d. random effect for transect nested within site and reef-depth
- $\varepsilon_{ijt}$ is the observation-level random effect.

Sites were the unit of sampling for the Gaussian random field. The spatial domain was discretised into a lattice mesh using Delaunay triangulation, with nodes defined by the observed locations, and the boundary was constrained using a shapefile of the GBR. A minimum node distance for the triangles was set to ensure that sites within reefs were in the same triangle (cut-off of 0.95/5 for inshore shallow and inshore deep models, and 0.95/20 for the offshore model). The mesh was then used to define the SPDE. Approximate posterior predictions of MAp were generated for each habitat at each reef from 1000 posterior draws to establish maintenance thresholds of MAp.

#### 1.2 Critical thresholds

The critical threshold for MAp was determined by fitting a Bayesian linear model (using the INLA package in R) to observed juvenile *Acropora* densities on inshore reefs. The aim was to estimate the MAp level at which juvenile *Acropora* densities drop below the amount needed to support sustained reef recovery, as defined for the critical threshold of the Juvenile Density indicator. The threshold targets inshore reefs because they have the highest macroalgal species richness, cover, and biomass, coinciding with increased turbidity, sedimentation, and nutrient levels (De'ath and Fabricius 2010, Ceccarelli et al. 2020, Fabricius et al. 2023).

The model included an estimate of the lag effect of MAp as a covariate. The MApLag covariate was calculated as the mean of MAp over the current and previous two years (where data were available). This averaging accounted for juvenile Acropora observations, including colonies up to 5 cm in maximum dimension, representing corals that have settled and grown over several years. Observations taken in the months following exposure to major floodwaters or cyclone damage were excluded from the averages, as these events can significantly reduce MAp for a short period (eq. 2).

Juvenile *Acropora* abundance was estimated using a Generalised Linear Regression with a negative binomial distribution and random effect for each site, nested within reef and depth. To obtain density estimates rather than abundance, an offset for the availability of settlement substrate, defined as the area occupied by algae as opposed to other benthic organisms or unconsolidated substrates (avail.area), was included.

${log(\mu_{ijk})}\sim\beta_{0}+ \beta_{1}\log\left( MApLab+0.1 \right)+ \upsilon_{{site}_{jk}}+ offset\left( avail.area \right)+ \varepsilon_{ijkt}$	$Y_{ijk} \sim NegBin(\mu_{ijk}, \theta)$	(equation 2)

Where:

- $\mu_{ijk}$ is the expected count of juvenile Acropora at transect *I*, site *j*, within reef-depth combination *k*,
- $\beta_{0}$ is the intercept
- $\beta_{1}log(MApLab+0.1)$ is the covariate for macroalgae prevalence (MAp)
- $\upsilon_{{site}_{jk}}$ is the independent and identically distributed (i.i.d.) random effect for site nested within reef and depth
- $offset\left( avail.area \right)$is the offset function for the area of available substrate
- $\varepsilon_{ijkt}$is the observation-level random effect.
- $\theta$ is the overdispersion parameter

The upper limit value of MAp (0.224) at which juvenile *Acropora* density reached the level required for rapid reef recovery (0.54 per m^2^, as established for the critical metric for the Juvenile Density indicator) was selected as the MAp critical threshold (Figure S2.1).

### 2 *Estimations of* observation metrics

A Bayesian hierarchical linear temporal MAp model was fit for each reef-depth combination. MAp was fit with a beta-binomial distribution, a binomial distribution and a binomial distribution with an observation-level random effect to help account for overdispersion, each with a logit link. The best-fit model was selected using WAIC. The model included an intercept, a population-level effect of report year (Year), and random effects for site (*k*, nested within reef/habitat) and transect (*j*, nested within site). Including ‘report year’ as a population-level effect maximises consistency in hindcasting as additional years are added to the model (eq. 3).

${logit(MAp}_{kjt}) \sim\beta_{0}+ \beta_{1}{year}_{t}+ \upsilon_{k}+ \upsilon_{jk}+ \varepsilon_{kjt}$ (equation 3)

Where:

- ${MAp}_{kjt}$is the expected macroalgae prevalence for each site-depth (k), transect (j) and year (t),
- $\beta_{0}$ is the intercept,
- ${year}_{t}$ is the fixed-effect of report year,
- $\upsilon_{k}$is an i.i.d. random effect for site-depth habitats,
- $\upsilon_{jk}$ is an i.i.d. random effect for transects nested within site-depth habitats,
- $\varepsilon_{kjt}$ is the observation-level random effect.

For each temporal reef/habitat model, estimates of MAp in each survey year were derived as 1000 draws from the posterior distribution.

### 3 Index Score Calculations

#### 3.1 Maintenance index score

Index scores were calculated using the posterior draws $\left( I_{(i,j)} \right)$ from the indicator model for a given year ($i$) and site-habitat ($j$). Each drawn posterior is paired with a drawn posterior from the maintenance model ($T_{m}$). The maintenance index score ($f(x)$) is calculated for each pair from the difference between observed values and reference threshold, distance (*x*; eq. 5), and scaled from 0-1 using a Cumulative Distribution Function of a logistic distribution for a given distance metric (eq. 4).

$f\left( x;0,1 \right)= \frac{1}{\left( 1+ e^{\left( -x \right)} \right)}$	(equation 4)

Where:

$x= {log}_{2}\left( \frac{I_{(i,j)}}{T_{m}} \right)$	(equation 5)

Using this approach, an MAp value equal to the maintenance threshold yields a score of 0.5. Scores below 0.5 were rescaled to extend the lower value of the scores to 0 when the temporal reef value of MAp was 0.8. Scores above 0.5 were not rescaled, as scores naturally range from 0 to 1 when MAp is 0.

#### 3.2 Critical index score

Index scores were calculated using the posterior draws $\left( I_{(i,j)} \right)$ from the indicator model for a given year ($i$) and reef-habitat ($j$). Each drawn posterior is paired with a drawn posterior from the critical threshold ($T_{c}$). The critical index score ($f(y)$) is calculated for each pair from the difference between observed values and reference threshold, distance (*y*; Eq. 7), and scaled from 0-1 using a Cumulative Distribution Function of a logistic distribution for a given distance metric (Eq. 6).

$f\left( y;0,1 \right)= \frac{1}{\left( 1+ e^{\left( -y \right)} \right)}$	(equation 6)

Where:

$y= {log}_{2}\left( \frac{I_{(i,j)}}{T_{c}} \right)$	(equation 7)

The MAp value that equated to a score of 1 was set to 0, and the critical MAp threshold (0.224; Fig. S2.1) established from the model described earlier was set to equate to a score of 0.5. Using this calculation, the distance metric naturally runs to a score of 1 when MAp is 0. In contrast to the maintenance calculation, distances below 0.5 were set to 0 in the critical score calculation.

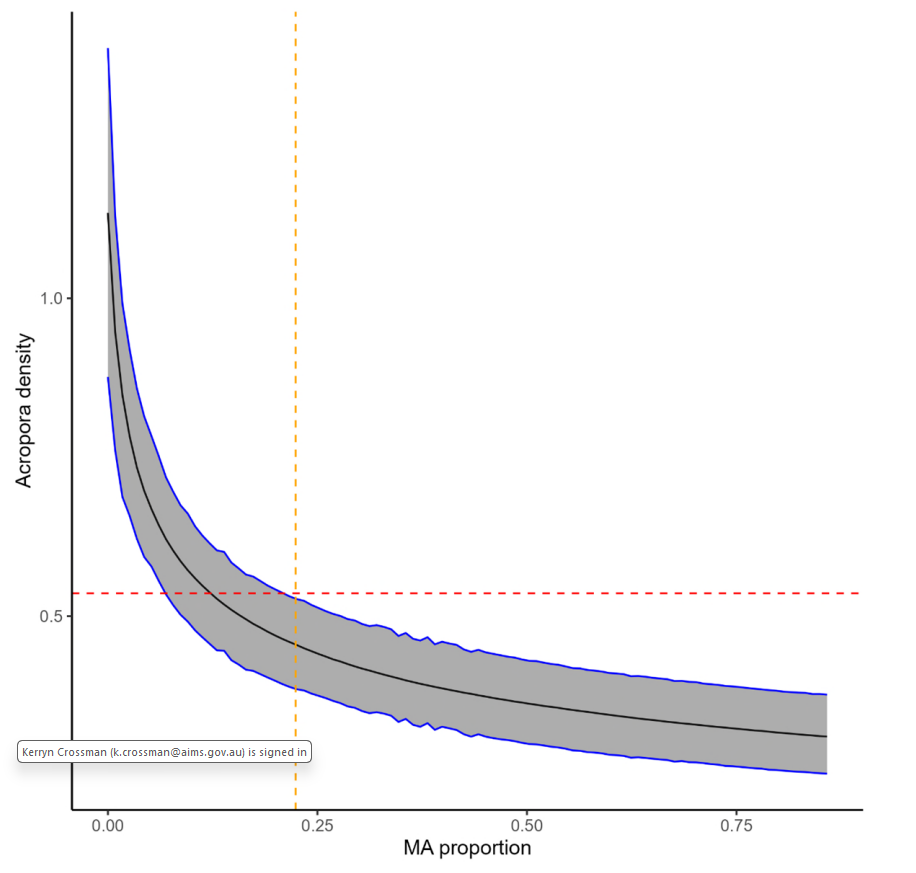

**Figure S2.1**: Predicted cover of juvenile *Acropora* density in response to MAp based on inshore reefs monitored by AIMS. Dashed lines indicate where the upper limit of MAp intersects with the level of juvenile *Acropora* density required for rapid reef recovery, as established for the critical metric of the juvenile coral indicator (0.54 per m2).
