## Supplementary material for "Beyond Coral Cover A framework for assessing the condition of coral reef habitats and informing conservation targets": ESM 4

ESM # 4: Community Composition Indicator

### 1 Estimation of thresholds

#### 1.1 Maintenance thresholds

The community composition maintenance metric measures the degree to which the composition of the coral community at a reef has deviated from community compositions previously observed at the reef during a reference period. The approach described here is a variant of the “local outlier factor” (LOF) methodology of multidimensional comparisons (Breunig et al. 2000), adjusted for ecological purposes. The reference period for each reef uses multidimensional benthic community data collected for all time points up to 2015. For reefs where initial surveys began after 2005, the first 10 observations serve as the reference period.

Cover of hard and soft coral taxa was converted to relative cover by square-root transforming raw proportions and then dividing each proportion by the sum of proportions for that survey. This transformation reduces the weight of abundant taxa relative to rarer taxa, making dissimilarity measures more sensitive to overall community change and not simply change in dominance.

The degree of deviation of the target observation from reference observations was measured using the Local Outlier Factor (LOF). LOF is a statistical value of dissimilarity generated from a multidimensional dataset. It compares a single data point (the target observation “T”; i.e., community composition of a reef in a given year) to the rest of the dataset by calculating the density of nearby neighbours in multidimensional space, and then contrasting this density with the density of its neighbours’ neighbours (secondary neighbours). The outlier factor has one parameter: “K”, the number of neighbours. For the maintenance metric, we used K=6. That is, the six observations from the reference period with the lowest Bray-Curtis dissimilarity to the target observation were selected as the reference neighbours, whose neighbourhood densities were compared to the target observation.

The Bray-Curtis distance between the target, *T,* and its K^th^ nearest neighbours (*K_D_*) was used to calculate the Reachability Distance (*RD*). The RD between *T* and each of its K^th^ nearest neighbours (*Xj*) is defined as the maximum of the k-distance of *Xj* and the Bray-Curtis distance between T and *Xj*. This process ensures that the RD reflects both the local density near *Xj* and the dissimilarity between T and *Xj* (Equation 1).

Eq. 1

$$Reachability Distance \left( T, Xj \right)=max(K_{D}\left( Xj \right), D\left( T,Xj \right))$$

Reachability Distances for the neighbours within K-distance of the target are converted into a Local Reachability Density (LRD) (Equation 2). In this equation, the Local Reachability Density of a target T is calculated as the inverse of the average distances D of T to all neighbours (X) within K-distance (*N_k_(T)*). This reflects the average distance at which T can be reached from its neighbours. In turn, LRD is calculated for the neighbours with K-distance, using secondary neighbours.

Eq. 2

$$Local Reachability Density \left( T \right)=\frac{1}{\left( \frac{\sum Xj\in N_{K}\left( T \right) RD(T,Xj)}{\left| N_{K}(T) \right|} \right)}$$

The Local Outlier Factor calculation then compares the LRD of the target “T” to the average LRD of its neighbours (Equation 3). The RD for each neighbour point is calculated, then converted to a set of RDs for each of the secondary neighbours (within the neighbour’s RD) (Equation 2), and then finally an LRD. These neighbour LRDs are then divided by the LRD of the target, creating an LRD ratio, before being averaged together by summing ratios and then dividing by the number of neighbours within RD of T.

Eq. 3

$$Local Outlier Factor \left( T \right)= \frac{\sum N\in N_{K}\left( T \right)\frac{{LRD}_{K}(N)}{{LRD}_{K}(T)}}{\left| N_{K}(T) \right|}$$

The resulting score is a Local Outlier Factor, a ratio of the average reachability of the target point to the average reachability of its six neighbours from the reference period. An LOF of 1 reflects a target that is surrounded by a similar density of neighbours compared to the density of secondary neighbours. LOF values increasingly greater than 1 reflect targets that are ‘outliers’ with low reachability relative to neighbours. This target is likely to be anomalous and deviate strongly from observations from the reference period.

### 2 Index Score Calculations

#### 2.1 Maintenance index score

LOF scores were standardised into indicator scores by using an inverse sigmoid function (Equation 4). This function corrected LOF values above 1 to scores below 1. Given that LOF scores are ratios (which are essentially log-transformed), this function uses the sigmoid scaling parameters (a and b) to adjust the scaling of the scores, such that an LOF value of 1.25 (i.e., target’s LRD is 80% of its neighbours) equated to a score of 0.5, and an LOF value of 2 (i.e., target’s LRD is 50% of its neighbours) equated to a score approximating 0.05. The sigmoid scaling parameters to achieve this transformation are a = 3 and b = 1.

Eq. 4

$$Indicator \left( T_{ab} \right)= \frac{1 / (1+ e^{aLOF-b})}{1 / (1+ e^{a-b})}$$

#### 2.2 Taxonomic changes

To investigate the contribution of each taxon to overall changes in community composition, mean relative abundance of each taxon was calculated over the baseline years (1995-2015), and the difference between the baseline mean and the relative abundance of each taxon in subsequent years was calculated to generate a table containing the annual difference in relative abundance of each taxon from the baseline mean.
