## Supplementary material for "Beyond Coral Cover A framework for assessing the condition of coral reef habitats and informing conservation targets": ESM 5

ESM # 5: Recovery rate Indicator

### 1 Estimation of thresholds

#### 1.1 Maintenance thresholds

The recovery rate maintenance metric is a measure of the magnitude of the difference between observed and expected hard coral cover during recovery from disturbance. Coral growth expectations were established at the habitat scale (deep and shallow) within bioregions. To achieve this, reef-level recovery trajectories from AIMS monitoring data up to 2020 were first parameterised using a two-phase logistic growth model (Warne et al. 2022). Data up to 2020 were required for adequate spatial coverage of reference trajectories across habitats within bioregions. The beginning of a recovery trajectory was recognised by a recorded disturbance in the database or by a 3% decline in the median of the posterior distribution of hard coral cover between consecutive observations at a reef. A minimum of four observations in a reference trajectory was required for model parameterisation.

The two-phase growth model consists of two generalised logistic growth models (Eq. 1), with a change point from the first model (first phase) to the second model (second phase) determined using a piecewise-defined population dynamic model (Murphy et al. 2022, Eq. 2)

The generalised logistic growth model is defined as:

$$\mathrm{Equation} 1: \frac{dC(t)}{dt} = \alpha C\left( t \right) \times\frac{1}{\gamma}\left[ 1-\left( \frac{C(t)}{K} \right)^{\gamma} \right] , t>t_{0}$$

Rate of change in cover

Competition for available space
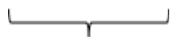

Coral growth

where K is the maximum % area the coral can cover, i.e., the available substrate, C(t) is the hard coral cover at time t (years), > 0 (1/years), α is the intrinsic rate of cover increase and γ > 0 is a non-dimensional generalisation parameter that controls the shape of the sigmoid growth curve (curve shape).

The piecewise-defined population dynamic model is:

$$Equation 2: \frac{dC(t)}{dt}= \left\{ \begin{aligned} f_{1}\left( C\left( t \right) \right) \mathrm{if} t\leq T, \\ f_{2}\left( C\left( t \right) \right) \mathrm{if} t\leq T, \end{aligned} \right.$$

where the rate of change in cover, switches from f1(C(t)) to f2(C(t)) at the change point t = T.

The two-phase recovery model required the recovery rate of the first phase to be lower than that of the second phase. To achieve this, we implemented the definition f1(C(t)) = αdf2(C(t)) where αd ∈ (0, 1) is the scale reduction factor, and f2(C(t)) corresponds to the generalised logistic growth model (Equation (1)). Further, we defined a duration, Td > 0, after which the recovery reverts back to Equation (1), and then substituted Equation (1) and initial exponential growth into Equation (2) to obtain the following two-phase recovery model definition:

$$Equation 3: \frac{dC(t)}{dt}= \left\{ \begin{aligned} \frac{\alpha_{d}\alpha}{\gamma}C\left( t \right)\left[ 1- \left( \frac{C(t)}{K} \right)^{\gamma} \right] \mathrm{if} t_{0}<t \leq t_{0}+T_{d} \\ \frac{\alpha}{\gamma}C\left( t \right) \left[ 1-\left( \frac{C(t)}{K} \right)^{\gamma} \right] \mathrm{if} t >t_{0}+T_{d} \end{aligned} \right.$$

Given an initial condition C(t0) = C0 and values for the parameters θ = [α, αd, γ, Td, K], an analytic solution is given by:

$$\mathrm{Equation} 4: C\left( t \right) \left\{ \begin{aligned} K\left\{ 1+ \left[ \left( \frac{K}{C_{0}} \right)^{\gamma}-1 \right]\exp\left( {-\alpha}_{d}\alpha\left( t-t_{0} \right) \right) \right\}^{-\frac{1}{\gamma}} , \mathrm{if}t_{0} <t \leq t_{0}+T_{d} , \\ K\left\{ 1+\left[ \left( \frac{K}{C_{0}} \right)^{\gamma}-1 \right] \exp\left( -\alpha\left( t-T_{d} \right) \right) \right\}^{-\frac{1}{\gamma}} , \mathrm{if}t>t_{0}+T_{d}, \end{aligned} \right.$$

To recognise the long-term but spatially variable presence of soft corals on reefs of the GBR, carrying capacity was conservatively reduced by 10% (the approximate temporal and spatial mean of soft coral cover in AIMS monitoring data). Carrying Capacity was also reduced by the cover of abiotic groups (sand and silt). That is,

𝐾 = 100 − 10 − abiotic cover

The two-phase model was parameterised in a Bayesian framework (Gelman et al. 2013) for each trajectory, yielding posterior densities for the four key parameters.

The four key parameters estimated by the model per trajectory were:

1) Growth rate - The intrinsic growth rate of the logistic growth curve in the second phase.

2) Scale reduction factor - The first phase growth rate as a proportion of the second phase growth rate.

3) Curve shape - The shape of the logistic curve, where values closer to 0 reflect Gompertz growth and values approaching 1 reflect logistic growth. Curve shape is assumed not to change between phases.

4) Change point - The time spent in the first phase in units of years.

Three criteria based on two-phase model parameters were identified as criteria that detected notably slow historical recovery trajectories: 1) Below the 15th percentile of the 2nd phase growth rate, 2) Duration of first phase >4 years & 1st phase growth rate < 0.04, 3) Duration of first phase >2 years & 1st phase growth rate <0.1. Trajectories meeting these criteria were deemed unsuitable for setting recovery rate expectations and were excluded from the reference trajectory pool.

For each reference trajectory, 2000 rows of growth rate, scale reduction factor, curve shape, and change point combinations were randomly sampled (500 from each of 4 chains), and the parameter rows were pooled across corresponding reefs within habitat and bioregion combinations. For 1000 random draws of rows (combinations of the 4 parameters) from the bioregional pool, the four model parameters were used, along with information about the amount of time that has passed since disturbance and the coral cover at, and time passed since, the previous survey, to predict expected hard coral cover for the current observation. This process yielded a posterior distribution of 1000 estimates of expected hard coral cover based on bioregional expectations of recovery rates.

### 2 *Estimations of* observation metrics

We used the posterior distributions of hard coral cover estimated from the temporal reef model developed for the coral cover indicator as the observed data.

### 3 Index Score Calculations

#### 3.1 Maintenance index score

The minimum and maximum levels of observed hard coral cover selected to set index scores of 0 and 1 were:

Score = 0, when *observed cover = expected cover/2*

Score = 1, when *observed cover =2 x expected cover*

An observed value equal to the expected value equated to a score of 0.5. Each posterior draw $I_{(i,j)}$ of observed hard coral cover for each reef-habitat (*i*) and year (*j*) from the temporal reef-habitat model was paired with a draw from the posterior distribution of expected hard coral cover from the two-phase model (*C*). The score was calculated as the difference (*x*) between each pair of observed and expected hard coral cover values (*I_(ij)_*; Eq. 5) and scaled to 0-1 using the cumulative distribution function of a logistic distribution for a given distance metric (Eq. 6).

$x= {log}_{2}\left( \frac{I_{(i,j)}}{C} \right)$	(equation 5)

$f\left( x;0,1 \right)= \frac{1}{\left( 1+ e^{\left( -x \right)} \right)}$	(equation 6)

In order to impose the minimum and maximum values for scores of 0 and 1 as described above, distances >1 were converted to 1, distances < -1 were converted to –1, and the values of the resulting distribution were rescaled between 0 and 1.
