## Supplementary material for "Beyond Coral Cover A framework for assessing the condition of coral reef habitats and informing conservation targets": ESM 6

ESM # 6: Coral Juvenile Density Indicator

### 1 Estimation of thresholds

#### 1.1 Maintenance thresholds

Maintenance thresholds were estimated as the average of historical observations (2005-2015) for each reef bioregion in the GBR (Fernandes et al. 2005, Kerrigan et al. 2010) and categorical survey depth (“shallow” and “deep”). For bioregions with small sample sizes across the monitoring datasets, they were grouped based on geomorphological, environmental, and ecological descriptions when appropriate.

We applied the Integrated Nested Laplace Approximation (INLA) framework, combined with the Stochastic Partial Differential Equation (SPDE) approach, to estimate the abundance of juvenile coral colonies (<5 cm in diameter). Abundance (*n_i_*) was modelled using a Poisson Generalised Linear Model (GLM) with a log link function, a spatially structured random effect ($\varphi_{s_{i}}$), that captures spatial autocorrelation, and an offset value representing the area (m^2^) of juvenile transects occupied by algae (including turf, coralline and macroalgae). This spatial random-effect component was approximated using a Gaussian Random Field (GRF) via the SPDE approach. To account for variability introduced by differences in survey methodology, temporal fluctuations, and site-specific effects, we included additional random effects for dataset, year, and site. These components help isolate spatial patterns from other sources of variation (see Eq. 1).

$\log\left( \mu_{{ijt}} \right)\sim\beta_{0}+ \varphi_{s_{i}}+ \upsilon_{i}+ \upsilon_{t}+ \upsilon_{j}+\varepsilon_{ijkt}$	(Equation 1)

Where:

- $\mu_{{ijt}}$is the expected juvenile abundance,
- $\beta_{0}$ is the intercept,
- $\varphi_{s_{i}}$ is the spatially structured random effect estimated through SPDE,
- $\upsilon_{i}$is an i.i.d. random effect for monitoring datasets,
- $\upsilon_{t}$ is an i.i.d. random effect for years,
- $\upsilon_{j}$ is an i.i.d. random effect for reef/habitat
- $\varepsilon_{ijt}$ is the observation-level random effect.

Sites were the unit of sampling for the Gaussian random field. The spatial domain was discretised into a lattice mesh using Delaunay triangulation, with nodes defined by the observed locations, and the boundary was constrained using a shapefile of the GBR. A minimum node distance for the triangles was set to ensure that sites within reefs were in the same triangle (cut-off of 0.95/20). The mesh was then used to define the SPDE.

To establish maintenance thresholds for total juvenile density at a bioregional level (per habitat), a hexagon grid was overlaid onto the sample area (hexagon width approximately 3km, an area approximately 564 hectares). Hexagon areas that sit over a reef were filtered, and the reef area within each hexagon was calculated. Approximate posterior predictions were generated for each hexagon from 1000 posterior draws. Posterior predictions at the bioregion level represented a weighted (proportional reef area) mean across the hexagons.

#### 1.2 Critical thresholds

The critical threshold for juvenile coral density was estimated as the minimum density of juvenile *Acropora* colonies required for sustained recovery to 30% hard coral cover, following disturbance, within 10 years. It was derived from Integral Population Projection Models (IPMs), in which different-sized coral colonies are modelled across a continuous spectrum of sizes and represented across different life-cycle stages (sensu Kayal et al. 2018, Cresswell et al. 2024). At each timestep, a coral may die (mortality) or transition to a different size, growing, shrinking (partial mortality), or staying the same size, with probabilities determined by Bayesian regressions of growth and survival (Figure S5.1). Reproduction was not included in the model because the objective of the exercise was to manipulate the number of juveniles that settled into the population.

Demographic rates for growth and survival were estimated from field observations across environmental gradients and major hard coral taxa using published and unpublished data sources (Doropoulos et al. 2015, Doropoulos et al. 2022, AIMS 2023, Madin et al. 2023 Hill and Hoogenboom, unpublished data; Hoogenboom, unpublished). We modelled yearly growth and survival rates of hard corals using Bayesian Generalised Linear Models (GLMs), accounting for size-dependent relationships that vary across geographic regions and local environmental conditions (sensu Cresswell et al. 2024).

Estimated demographic rates were then used to parameterise IPMs and to predict coral cover change over time across taxa and habitats (Inshore and Offshore), as the abundance of juvenile corals recruited into the population each year varied. Each simulation was repeated 4,000 times to estimate the variability in population growth responses across scenarios.

Necessary model assumptions to support our objective included recovery initiating from 1% coral cover, open populations, no settlement space or carrying capacity limitations, *Acropora* as the primary coral genus driving recovery and a modelled reef habitat of 100 m^2^_._

Fixing the densities of other major coral taxa to 0.6 ind.m^-2^, we calculated the total coral cover at 10 years when varying the density of *Acropora* juveniles arriving each year into the population in 0.2 ind.m^-2^ increments. Model predictions were evaluated using numerical optimisation algorithms (Nash 1990) to assess the minimum yearly density of juvenile *Acropora* colonies required on a given inshore or offshore reef to ensure 30% total coral cover in 10 years following acute disturbances.

For inshore reef habitats, the minimum density of *Acropora* juveniles required to be observed yearly to ensure recovery to 30% total coral cover in 10 years was estimated at 0.54 ind.m^-2^ (0.49 – 0.59 ind.m^-2^, 95% credible intervals; Fig. S5.2). To achieve the same outcome on offshore reefs, the estimated minimum *Acropora* juvenile density required yearly was 0.95 ind.m^-2^ (0.90 – 1.00 ind.m^-2^, 95% credible intervals; Fig. S5.2).

### 2 *Estimations of* observation metrics

Juvenile abundance was estimated using a Generalised Linear Regression with a negative binomial distribution and random effect for each site, nested within reef and depth. To obtain density estimates rather than abundance, an offset for the availability of settlement substrate, defined as the area occupied by algae as opposed to other benthic organisms or unconsolidated substrates (avail.area), was included.

${log(\mu_{ijk})}\sim\beta_{0}+ \beta_{1}\log\left( MApLab+0.1 \right)+ \upsilon_{{site}_{jk}}+ offset\left( avail.area \right)+ \varepsilon_{ijkt}$	$Y_{ijk} \sim NegBin(\mu_{ijk}, \theta)$	(Equation 2)

Where:

- $\mu_{ijk}$ is the expected count of juvenile at transect *I*, site *j*, within reef-depth combination *k*,
- $\beta_{0}$ is the intercept
- $\beta_{1}log(MApLab+0.1)$ is the covariate for macroalgae prevalence (MAp)
- $\upsilon_{{site}_{jk}}$ is the independent and identically distributed (i.i.d.) random effect for site nested within reef and depth
- $offset\left( avail.area \right)$is the offset function for the area of available substrate
- $\varepsilon_{ijkt}$is the observation-level random effect.
- $\theta$ is the overdispersion parameter

For each temporal reef-habitat model, estimates of juvenile density in each survey year were derived as 1000 draws from the posterior distribution

### 3 Index Score Calculations

#### 3.1 Maintenance index score

Index scores were calculated using the posterior draws $\left( I_{(i,j)} \right)$ from the indicator model for a given year ($i$) and reef-habitat ($j$). Each drawn posterior is paired with a drawn posterior from the maintenance model ($T_{m}$). The maintenance index score ($f(x)$) is calculated for each pair from the difference between observed values and reference threshold, distance (*x*; Eq. 4), and scaled from 0-1 using a Cumulative Distribution Function of a logistic distribution for a given distance metric (Eq. 3).

$f\left( x;0,1 \right)= \frac{1}{\left( 1+ e^{\left( -x \right)} \right)}$	(Equation 3)

Where:

$x= {log}_{2}\left( \frac{I_{(i,j)}}{T_{m}} \right)$	(Equation 4)

A minimum and maximum juvenile density value that equated to scores of 0 and 1, respectively, were selected separately for inshore deep, inshore shallow and offshore deep habitats as there was a clear distinction at this spatial scale in maintenance threshold values. For each habitat, the minimum value equating to a score of 0 was a juvenile density of 0. The maximum value was set as double the maximum baseline value for the habitat. The resulting maximum values of juvenile density equating to a score of 1 were 8, 18 and 34 for inshore shallow, inshore deep and offshore deep habitats, respectively.

Using this approach, a juvenile density equal to the maintenance threshold yielded a score of 0.5. Scores above 0.5 were rescaled to extend the upper value of the scores to 1 when the temporal reef value of juvenile density was either 8,18, or 34, depending on the corresponding habitat cap. Scores below 0.5 were not rescaled, as scores naturally approach 0 when juvenile density is 0.

#### 3.2 Critical Index score

Index scores were calculated using the posterior draws $\left( I_{(i,j)} \right)$ from the indicator model for a given year ($i$) and reef-habitat ($j$). Each drawn posterior is paired with a drawn posterior from the critical threshold ($T_{c}$). The critical index score ($f(y)$) is calculated for each pair from the difference between observed values and reference threshold, distance (*y*; Eq. 6), and scaled from 0-1 using a Cumulative Distribution Function of a logistic distribution for a given distance metric (Eq. 5).

$f\left( y;0,1 \right)= \frac{1}{\left( 1+ e^{\left( -y \right)} \right)}$	(Equation 5)

Where:

$y= {log}_{2}\left( \frac{I_{(i,j)}}{T_{c}} \right)$	(Equation 6)

The thresholds of juvenile *Acropora* spp. The density required for fast coral recovery on inshore and offshore reefs (0.54 and 0.95, respectively) established from the IPMs described earlier was used to set the 0.5 score of the critical index. A maximum *Acropora* spp. density value that equated to a score of 1 was set to four times the critical threshold, so that values of 2.16 and 3.8 equated to scores of 1 for inshore and offshore reefs, respectively.

Scores above 0.5 were rescaled to extend the upper value of the scores to 1 when the temporal reef value of juvenile *Acropora* density reached the corresponding value set as the maximum cap. Distances below 0.5 were converted to 0.

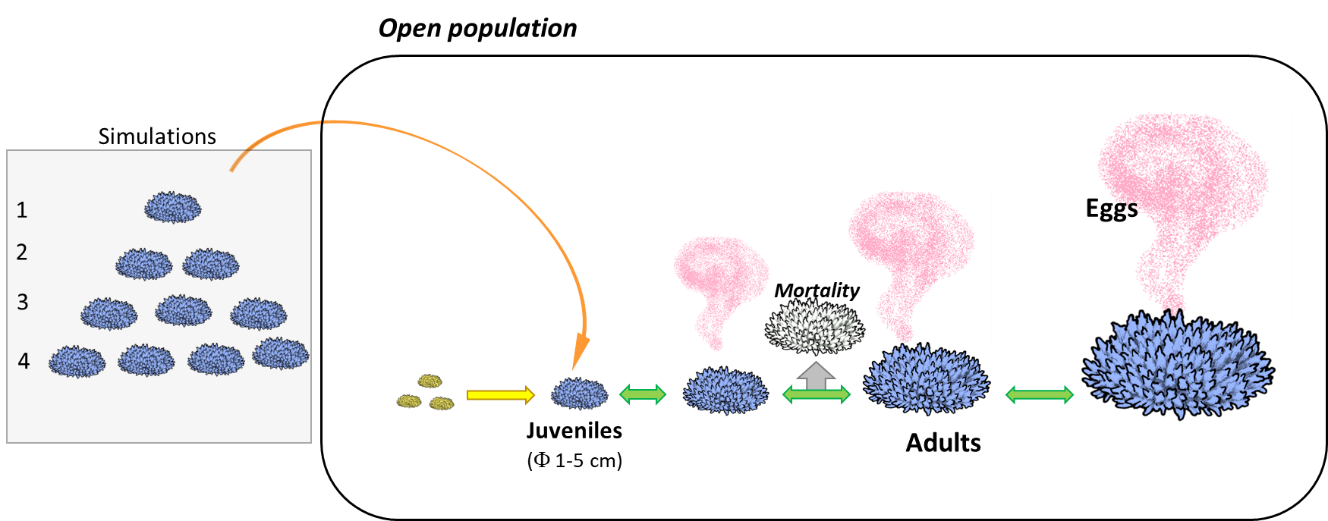

**Figure S5.1.** A schematic representation of the coral life cycle and the main modelled processes. Different-sized coral colonies are modelled across a continuous spectrum of sizes and represented in the life cycle. At each timestep, a coral may die (mortality) or transition to a different size (green arrows), growing, shrinking (partial mortality), or staying the same size. Mature colonies may reproduce once per timestep, releasing gametes into the water column (pink), which may fertilise and develop into larvae. Here, we simulated an open population in which the number of juveniles settling into the population was manipulated to investigate the minimum number required to maintain rapid recovery of coral cover.

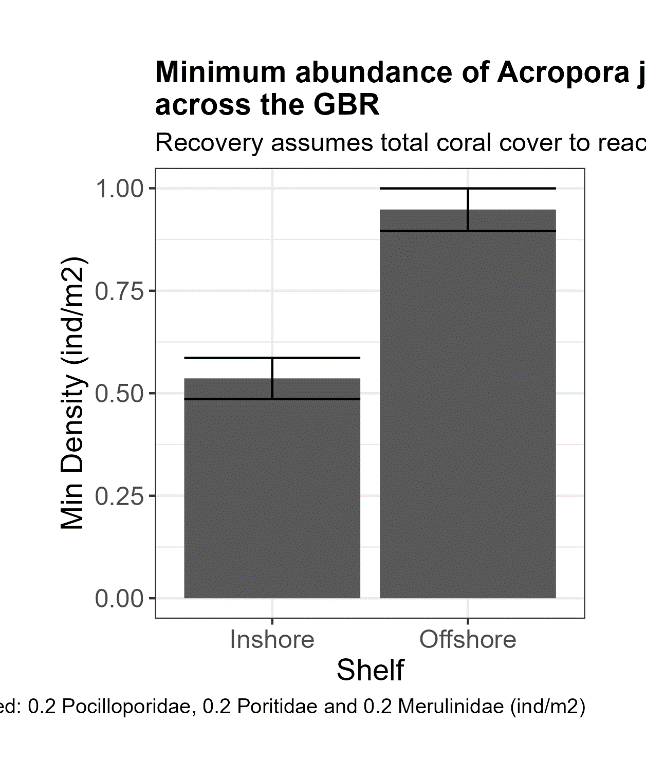

**Figure S5.3.** Minimum density of *Acropora* juvenile corals estimated for recovering communities to reach 30% total coral cover in 10 years after a severe disturbance. The figure shows the mean and confidence intervals (95%) of *Acropora* densities across shelf positions.
